# A reactant-dependent algebraic framework to define, construct, characterize and assess the redundant forms of a biochemical network

**DOI:** 10.1101/2024.03.07.583960

**Authors:** Siddhartha Kundu

## Abstract

The intracellular milieu, replete with concomitantly occurring biochemical networks, constitutes a complex physicochemical environment wherein macromolecules interact and whose activities can be modified by small molecules. These metabolism-contributing, signal-transducing and response-regulating networks, and the redundant forms that result, thereof, are responsible for complex biochemical-, physiological- and clinical-outcomes. It is unclear whether a unique mathematical structure, which is concurrently resolvable into redundant, unequal and functionally relevant substructures, can be ascribed to a biochemical network. Here, we develop a reactant-dependent algebraic framework to define, construct, characterize and assess the redundant forms of a biochemical network. We deploy a constraint-based approach to populate and select plausible reaction vectors in accordance with known paradigms of biochemically relevant outcomes. There is a strict lower bound for the number of participating reactants with integral changes, across all molecules of a reactant, which are Boolean (zero, signed non-zero) and combinatorially distributed across linearly independent plausible reaction vectors. The reactants and the selected plausible reaction vectors form a set of unique stoichiometry number matrices, each of which represents a distinct biochemical network which when multiplied by a set of positive and non-unitary rational numbers will generate a library of equivalent rational number matrices. The matrices that comprise this library are unequal, disjoint, form a semigroup with respect to addition, share the same null space and represent the redundant forms of a biochemical network. The presented framework is theoretically sound, mathematically rigorous, readily implementable, easily parameterized and can be used to assess the biomedical relevance for the redundant forms of a biochemical network. We conclude our thesis with plausible explanations into the genesis of formaldehyde resistance and thence nosocomial infections by *Klebsiella spp*.

## Introduction

Biochemical networks of enzyme-mediated transformations, exchange- and diffusion-reactions, and formation/disassembly of molecular complexes) are regulated by short (feedback mechanisms, post-translational modifications)- and long (*de novo* synthesis, degradation)-term mechanisms [1, 2]. A biochemical network is either studied by optimizing the biomass of a reactant of interest and enumerating elementary vectors or is data-driven, inverse and inferential. Algorithms which optimize one or more reactants are likely to be biased in choice, type and usage of constraints, and will result in studies with reduced biomedical relevance despite being numerically sound [1, 3–6]. Enumeration-based algorithms on the other hand, involve extensive computations that are NP-hard and will become intractable for large biochemical networks [2, 6–8]. The availability of- and accessibility to-empirical- and clinical-datasets has the potential to recreate, without bias, relevant data-driven biochemical networks, infer directionality for a given biochemical network (polynomial equation) or represent an arbitrary state of a modelled network (continuous-time Lyapunov equation) [9–12]. In these approaches perturbations are modelled as stochastic fluctuations and are used to derive the covariance (weak)- or correlation (strong)-interaction matrices for the set of nodes that comprise the modelled biochemical network [13–15].. This reliance on empirical data both, *a priori* and *a posteriori*, along with the computational complexity in computing the Jacobian are limitations in the inferential study of biochemical networks [13–15].

These limitations notwithstanding, inferred biochemical networks have a role in precision medicine and data mining [16–20]. For example, the use of guidance-based radio- and immune-therapeutics in managing chronic inflammatory diseases, neurodegenerative pathologies and recalcitrant and recurring malignancies is being used to assess dose response and predict outcome such as the propensity, progress and resolution of an adverse drug reaction [16–20]. In particular, these investigations combine the expression/abundance profiles of system-specific panels of biomarkers (genes, proteins, small molecules) into one of several machine learning (ML)-algorithms and predict patient response and/or compliance [18–20]. However, despite these advancements the primary or raw data (time-series kinetic data, “omics”-based datasets) is dependent on the platform used, experimental design and will require several numerical assumptions and extensive- and varied-normalization protocols [9–14]. Additionally, the use of expensive equipment, reagents and manpower can potentially restrict the utility of these investigations. In contrast to the aforementioned discussion on large scale modelling and analysis, causal networks and the inferences that result thereof, examine the effects of variable stimuli on several equally plausible and well-defined outcomes of a few nodes. Although causal network modelling has had several proven successes in investigating discrete real world events, the requirement for reference data, simple architecture and simpler rules, precludes consideration of complex biochemical systems other than for mining large biological datasets for lists of statistically significant genes/proteins of well characterized networks and in response to defined perturbations [21–25].

As opposed to the data-driven approach discussed vide supra, algebraic models for biochemical networks are usually formulated as population models [26, 27]. In particular, chemical reaction networks, with regards to the partaking reactants, are modelled as either self-sustaining/persistent once initialized or limiting (zero-occurring) [28, 29]. Although these investigations have enhanced our repertoire of modelling strategies for functional biochemical networks, novel data-analytics and parameter-free frameworks, the absence of biochemically-relevant parameterization may mitigate the potential biomedical impact and relevance of these studies [30, 31]. The dissociation (Kd)-, association (Ka)- and Michaelis-Menten (Km)-constants are important parameters in comprehending the outcome of a biochemical reaction (non-enzyme, enzyme) and are empirically determined (dual polarization interferometry, surface plasmon resonance, stopped-flow spectroscopy), inferred (regularized least squares pKa spectrum, static binding isotherms) and/or theoretically derived [32–38]. The probable dissociation constant, for the i^th^-reaction, is an unbiased numerical measure that is computed from a null space-generated subspace of combinatorially summed non-redundant and non-trivial vectors of the stoichiometry number matrix for a biochemical network [7, 8]. The probable dissociation constant is a strictly positive real number which can unambiguously compute the forward-, reverse- or equivalent-outcomes for a reaction [7, 8].

A biochemical network is modelled as an undirected graph with a finite number of nodes and edges, as opposed to a specific pathway which is, in contrast, rule-based and a directed version of the same. However, modelling the multitude of genome-scale derived data points mandates the use of vertex-bounded polytopes such as an n-dimensional simplicial complex with n + 1 affinely-independent vertices/nodes and a finite number 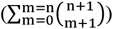 of faces. Here too, the absence of a biochemical rationale for the all- or none-manner in which the architecture of a network is perturbed whence node(s) are added/subtracted, choice of polytope, computational intractability at higher dimensions and the number of steady-state mapping constraints present unique challenges [39–41]. Redundant forms of a biochemical network (molecular, modular, network-level), are not uncommon, contribute significantly to physiological-, cellular- and biochemical-homeostasis (complement activation, blood clotting pathways, circadian rhythms) and are well-characterized (KEGG, MetaCyc) [42–46]. Relevance-studies, for these forms, include node-deletion (knock-out) or -inclusion (knock-in) with sensitivity analysis (single nucleotide polymorphism (SNP)-mapping) and last common ancestor (LCA)-tracing of molecular variants for epidemiological studies (source of an infecting pathogen, pedigree analysis, phylogenetic analysis) [3–7, 47, 48]. It is however, unclear whether a unique mathematical structure, which is concurrently resolvable into redundant, unequal and functionally relevant substructures, can be ascribed to a whole biochemical network. This paucity of mathematical descriptors notwithstanding, network-level redundant forms of a biochemical network occur in groups of organisms with significant taxonomic relatedness (species, strains, genotypes), are likely to facilitate herd immunity and may confer-/transfer-resistance to infectious- and chemotherapeutic-agents [49].

Here, we seek network-level redundant forms of a biochemical network, which are unequal, have a sound theoretical basis, mathematically rigorous, biochemically relevant and amenable to minimal perturbation(s). In order to accomplish this, we will develop a reactant-dependent algebraic framework to define and construct the redundant forms of a biochemical network, study its properties via suitable metrics (mathematical, computational) and assess biomedical relevance of the same (known biochemical pathways). In section 2, pertinent definitions and concepts in the form of biochemically relevant Constraints will be introduced and discussed. In “Methods” (section 3), we will describe the mathematical and computational tools needed to complement the theoretical assertions that have been outlined. This section will also describe the derivation of network-specific metrics which can be used to parameterize and establish relevance of the developed algebraic framework. In section 4 (“Results”), a detailed examination of the mathematical properties of the developed framework will be presented along with in-depth computational studies of known biochemical pathways. The “Discussion” (section 5) will highlight the rationale, motivation and biomedical relevance of this framework. The study concludes (section 6) by summarizing the salient features, major findings and limitations of this study. Detailed proofs, in order of appearance, for the mathematical formalism deployed, i.e., Constraints (Cs), definitions-environment (D), lemmas (L), theorems (T), corollaries (C) and propositions (P), are included in section 7 as supplementary material (STx) and cited wherever applicable

## 2 Definitions, concepts and preliminary results for a reactant-specific algebraic framework

Biochemical networks comprise interactions between finite numbers of molecular species of distinct reactants. The “reactions” that ensue are parameterized using enzyme-mediated catalysis (Km, Kcat, turnover number, catalytic efficiency), membrane-limited transport and macromolecular complex formation with disassembly (Kd, Ka).

### 2.1 Dissociation constants as units of change for reactants of a biochemical network

Consider a bimolecular non-enzymatic reaction,

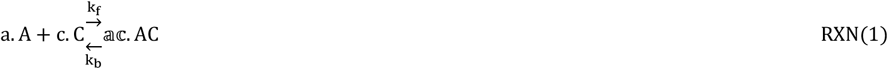

where,

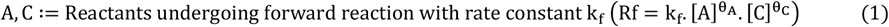

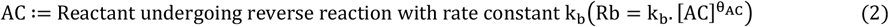

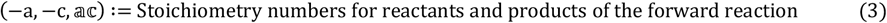

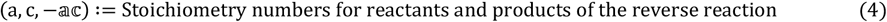

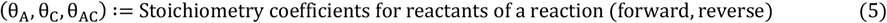

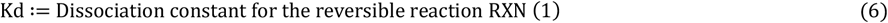

Let us assume that this reaction takes place at a steady state,

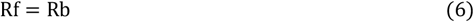

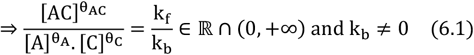

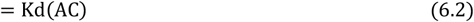

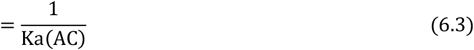

We can extend the annotation viz. **Eqs. (1–6)** to define the forward and reverse reactions,

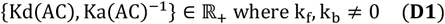

Let us describe a stoichiometry number for a reactant(s) from the dissociation constant with the linear map (h),

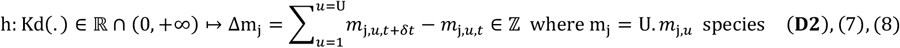

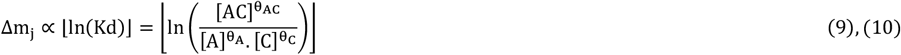

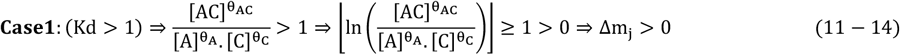

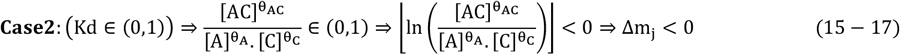

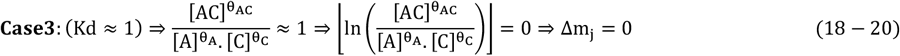

This map is onto and many-one since,

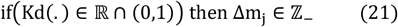

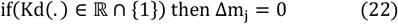

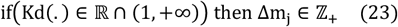

We now describe a biochemical network of j-indexed (j = 1, …, J) m-reactants (m_1_, …, m_J_) where each m_j_ is comprised of U. *m*_j,*u*_ species,

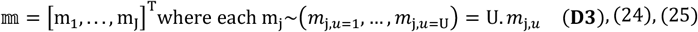

The reactions that comprise this network are i-indexed (i = 1, …, I) r-reactions (r_1_, …, r_I_) each of which is represented by a dissociation constant which we can rewrite as,

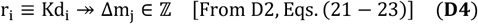

We will express the stoichiometry number for each reactant in terms of a reaction-specific dissociation constant in accordance with **D2-D4**. Since we are concerned with biochemical networks within a cell we will subsume a reactant’s contribution to each and every reaction, i.e., a reaction vector (RV),

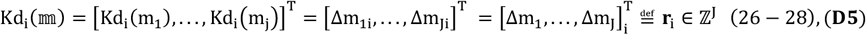

### 2.2 Constraints to construct a reactant-dependent algebraic framework

The intracellular milieu is an interplay of physico- and bio-chemical processes working in concert to achieve complex function. In our study we define and classify known instances of redundancies that are observed for a biochemical network in terms of the number of reactants. In order to assign a numerical measure to this constraint we will populate each RV in accordance with known biochemical paradigms.

#### 2.2.1 Constraints to define, describe and populate a plausible reaction vector

The basic unit of our algebraic framework is the plausible reaction vector (PRV) which unlike an RV will be, once populated, numerically tenable and representative of a true biochemical reaction.

##### Constraint 1

**(Cs1):** The contribution to the i^th^-PRV by J-reactants is Boolean (non-zero, zero),

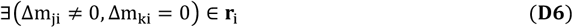

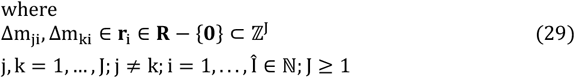

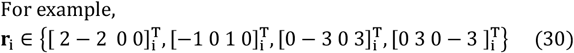

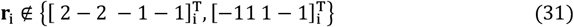

##### Constraint 2

**(Cs2)**: The cardinality of a set of i-indexed 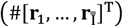, for a biochemical network with j-indexed J-reactants (Δm_1_, …, Δm_J_), is a function of the number of unique reactants that partake in the network.

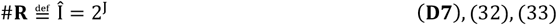

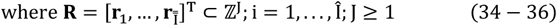

In order to uphold the premise of this study, i.e., a framework to model and characterize the redundant forms of a biochemical network, we will exclude PRVs that contribute to molecular (isoforms, isoenzymes, alternate splicing, pseudogenes, duplicate genes, truncated proteins)- and modular (futile cycles, sequestrations) –redundancy, for a biochemical network.

##### Constraint 3

**(Cs3)**: The PRVs used to construct the algebraic framework must be linearly independent,

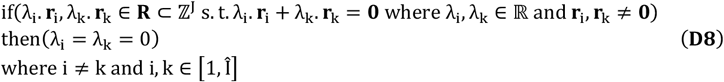

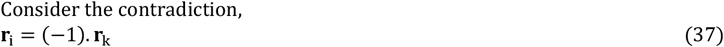

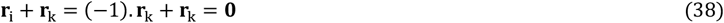

We will ensure that **Cs3** remains relevant at all stages of the modelling process by using a combinatorial approach to select- and populate-PRVs and identify reactant classes. The latter are combinations of subsets of reactants taken simultaneously. Additionally, we will preclude PRVs that are unlikely to be relevant from a biochemical perspective. For example, a zero vector means that every reactant is non-contributing, which is improbable given the complexity of the intracellular milieu (Δm_ji_ = 0 ∀ j; **D9, Eq**. (**39**)). It is equally improbable that every reactant will participate in every reaction (futile cycles, sequestration by membrane-bound compartments, feedback loops, partial reactions, competing metabolites) (Δm_ji_ ≠ 0 ∀j; **D10, Eq**. (**40**)). On the other hand, a singly signed non-zero entry-populated PRV of the form {Δm_ji_, Δ m_ki_} where, 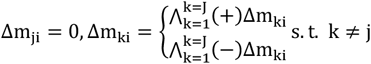 and k = 1, …, J; **D11, Eq**. (**41**) will indicate partial- or incomplete-reactions for a reactant or a product.

##### Constraint 4

(**Cs4**): The cardinality for the subset of excluded PRVs 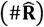, for J-reactants, is

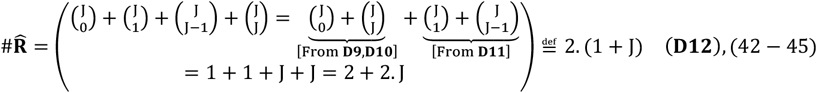

##### Constraint 5

(**A5**): The revised cardinality for the set of J-dependent PRVs,

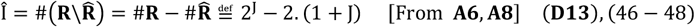

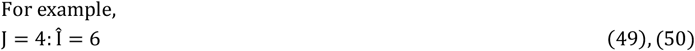

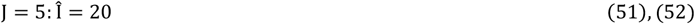

We will utilize signed non-zero reactants ((+)Δm_ji_, (−)Δm_ki_ where k ≠ j, k = 1, …, J) to populate a PRV in terms of their role in a reaction. A molecular species that is utilized/converted by a biochemical reaction is a reactant and is annotated as ((−)Δm_ji_) **(D14**). A product is defined as a molecular species which is formed by a biochemical reaction ((+)Δm_ji_) **(D15**). Clearly, **D14** and **D15** are only possible if the generating reaction is strictly forward or reverse (Kd_i_ ∈ ℝ_+_ − {1}). In contrast, a reactant without a net contribution (Δm_ji_ = 0) is likely to be due to uncertainty in the raw data or even an equivalent reaction (Kd_i_ ∈ ℝ_+_ ∩ {1}) [7, 8].

##### Constraint 6

**(A6)**: The stoichiometry contribution by any j-indexed m-reactant (Δm_j_) to a PRV is [**From D2, D3, D14, D15**],

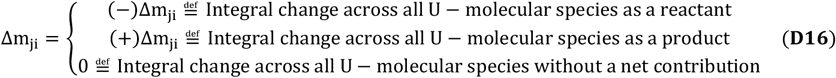

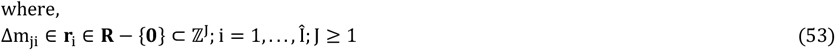

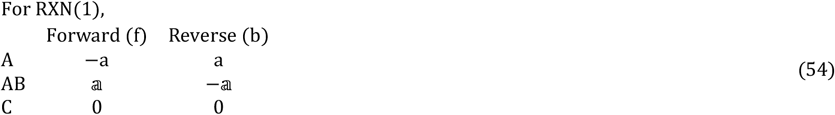

##### Constraint 7

**(Cs7)**: The i^th^-PRV must have at least one pair of reactants that are differently/oppositely signed,

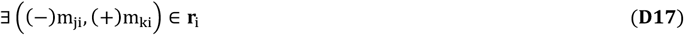

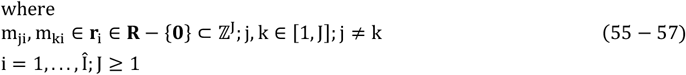

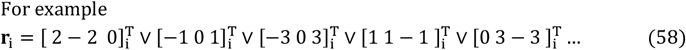

##### Constraint 8

**(Cs8):** The i^th^-PRV must comprise two non-zero and two zero entries/reactants,

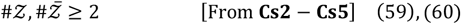

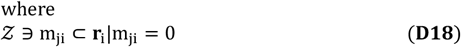

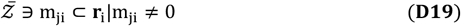

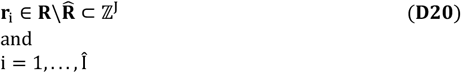

##### Constraint 9

(**Cs9**): The lower bound for the cardinality of an i^th^-PRV is,

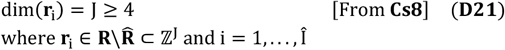

On the basis of **Cs1-Cs9** we will now refine our definition for a PRV as a non-zero, linearly independent vector with differently signed stoichiometry numbers for contributing reactants and zero values for non-contributing members both of which will not be less than 2 entries. We can now formally define the lower bound for the number of reactants (min(J) ≥ 4; **D22** [From **Cs9])** that will result in network-level redundant forms of a biochemical network thereby distinguishing it from molecular (J = 1)- and modular (J = 2,3)-versions of the same.

#### 2.2.3 Constraints to assemble a reactant-dependent stoichiometry number matrix

We have defined the criteria to populate and select subsets from the PRVs. We now describe Constraints that will combine selected PRVs into unique reactant-specific stoichiometry number matrices.

##### Constraint 10

(**Cs10**): The number of distinct subset(s) that J-reactants is divided into, where each subset comprises J ∈ [s + 1, J − 2] reactants that are concurrently considered, is computed from an s-indexed 𝕁-membered ordered list,

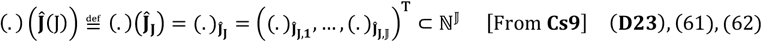

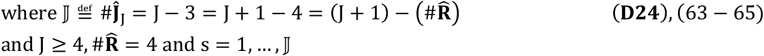

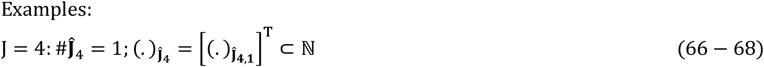

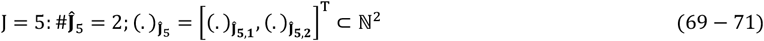

##### Constraint 11

(**Cs11**): We denote 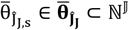 as the subset of a J-reactant dependent biochemical network where 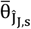 is the number of reactants considered simultaneously,

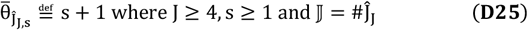

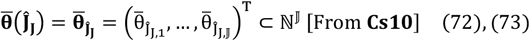

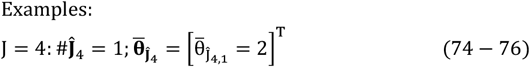

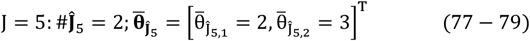

##### Constraint 12

**(Cs12):** We denote 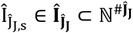 as the subset of a J-reactant dependent biochemical network where 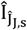 is the number of selected PRVs,

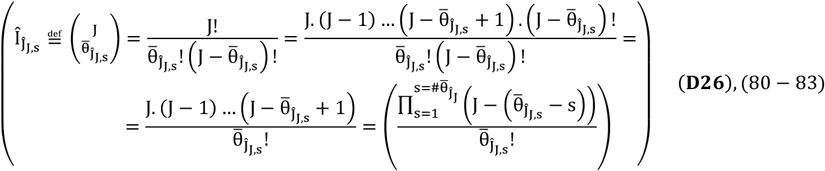

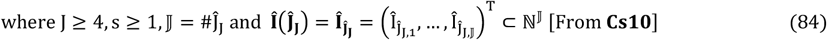

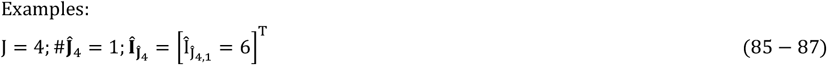

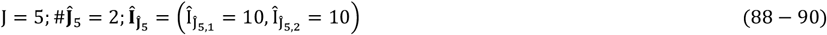

##### Constraint 13

(**Cs13**): A subset of i-indexed **r-**PRVs 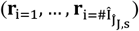 will form a stoichiometry number matrix 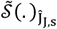,

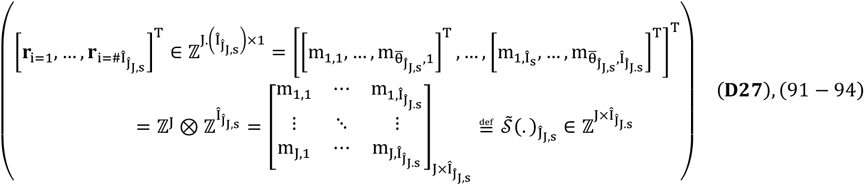

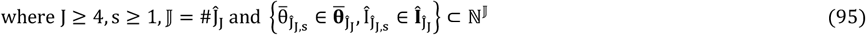

##### Constraint 14

(**Cs14**): The rank of 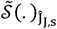 corresponds to the number of 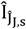-PRVs,

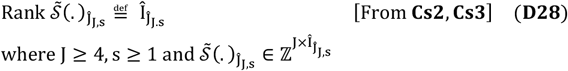

Whilst **Cs10-Cs14** result in a generic matrix 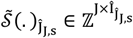 where s ≥ 1, the choice of reactants (s + 1, non-zero, 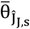; combinatorial, without replacement) in tandem with the corresponding 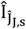-PRVs will result in different outcomes or stoichiometry number matrices.

##### Constraint 15

**(Cs15):** We denote 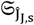 as a class-specific subset of *x*-indexed 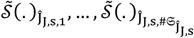-distinct stoichiometry number matrices for a J-reactant dependent biochemical network with 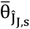-reactants taken concurrently, without replacement and in tandem with 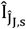-PRVs,

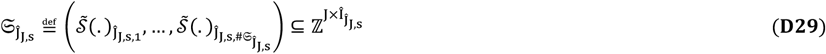

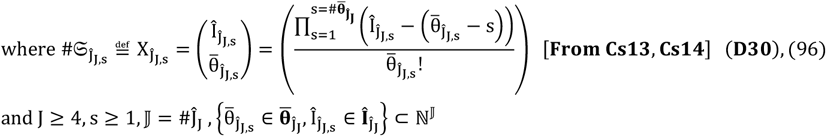

In order to explore the definitions discussed thus far, we present several numerical case studies for greater clarity (**Table 1**). We now present some preliminary results on the basis of **Cs1-Cs15**.

**Table 1:**
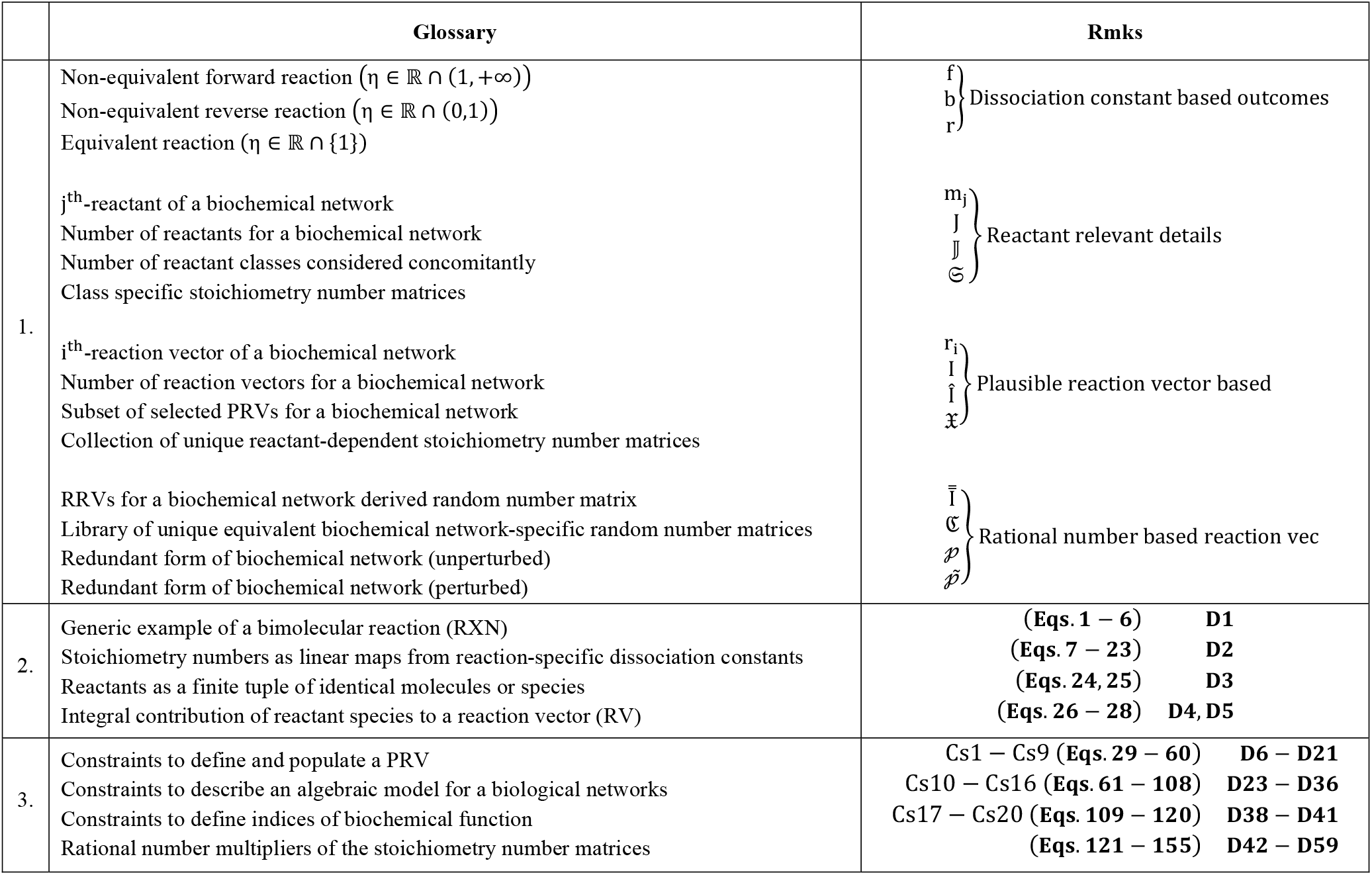
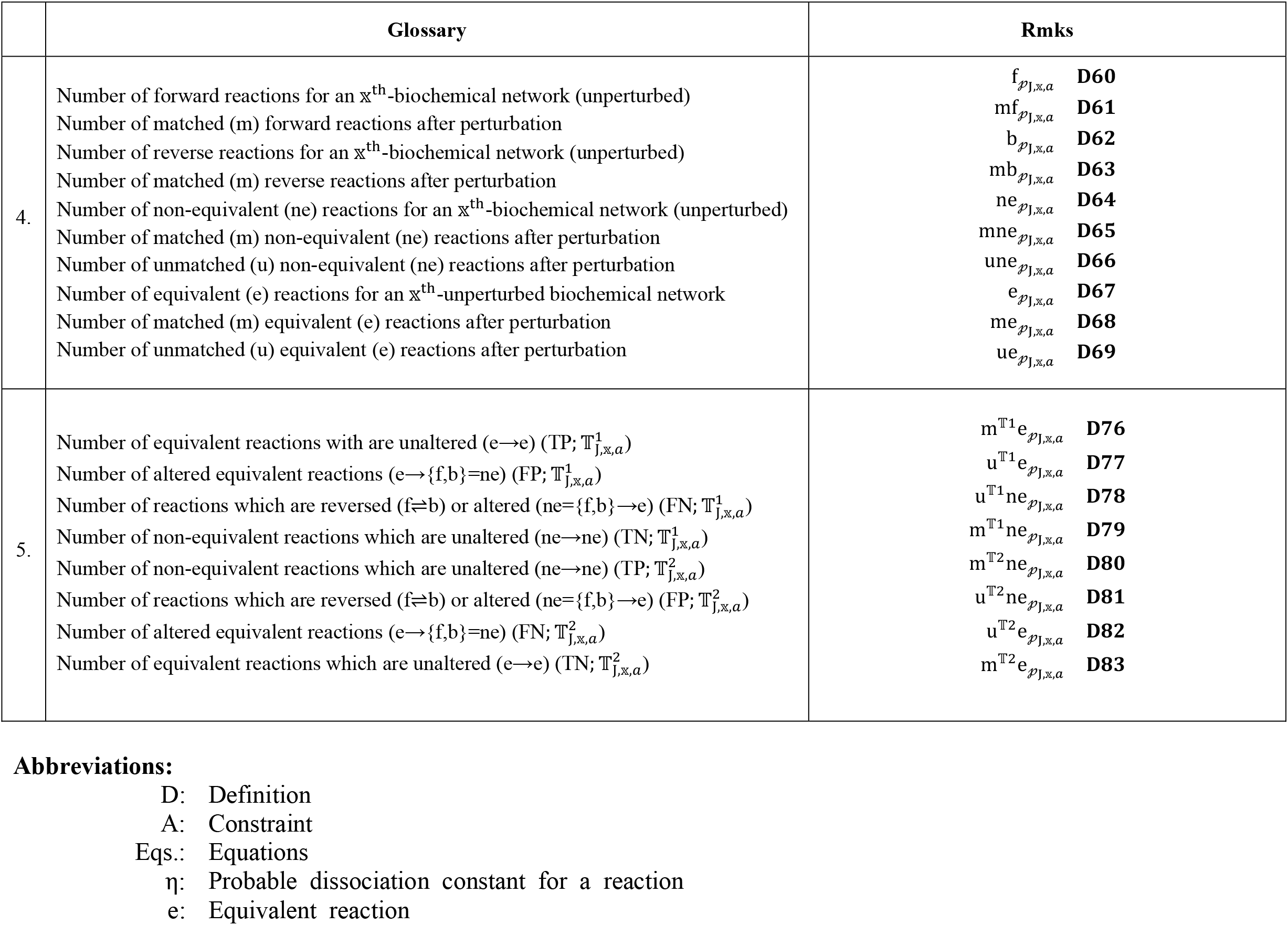
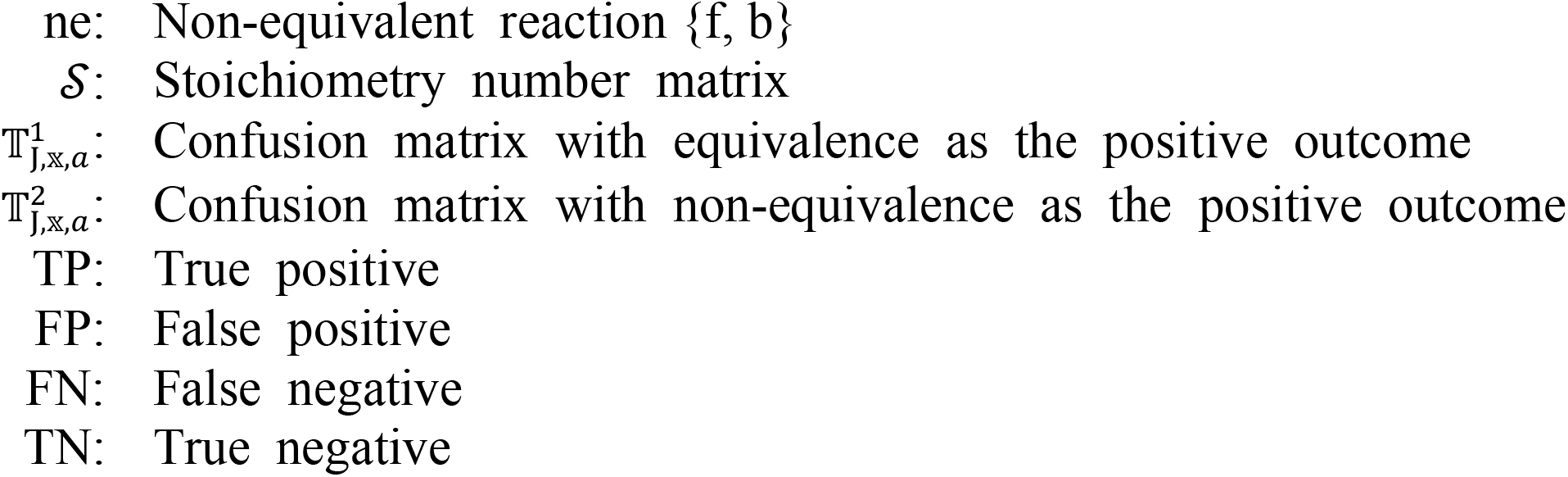
Glossary to define, construct, describe and analyse an algebraic framework to model and characterize redundant forms of a biochemical network.

##### Theorem 1

(**T1; STx1: Eqs. (1–7)**): There exist *x*-indexed 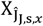-membered subset(s) of stoichiometry number matrices for a J-dependent biochemical network 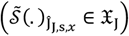 and for 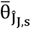-reactants (concurrent, without replacement) and in tandem with 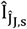-PRVs,

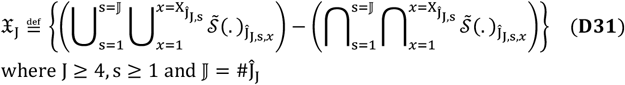

##### Corollary 1

(**C1; without proof**): The stoichiometry number matrices that comprise the superset of J-dependent biochemical networks 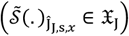 are non-overlapping, unique and unequal (From **T1**),

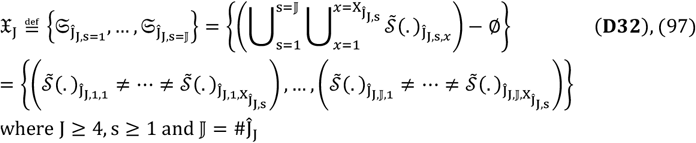

##### Corollary 2

(**C2**; **without proof)**: A J-reactant dependent superset 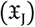 can be expressed as a vector of the independent classes of stoichiometry number matrices that comprises it 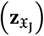,

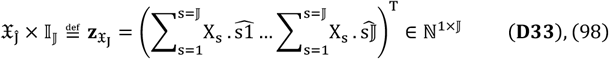

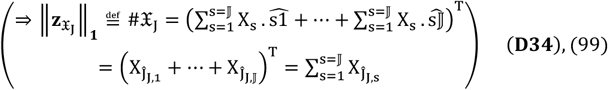

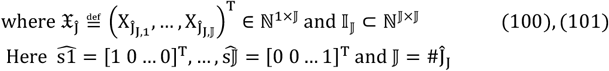

##### Theorem 2

(**T2**; **without proof)**: The J-reactant dependent superset 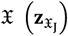 of stoichiometry number matrices can be expressed as a unit vector 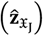,

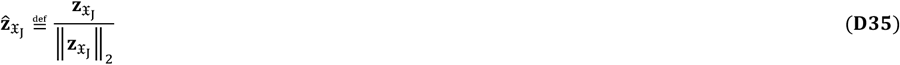

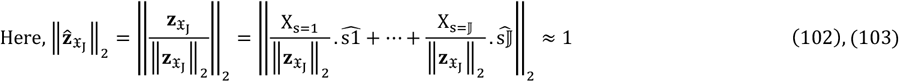

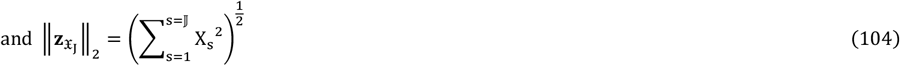

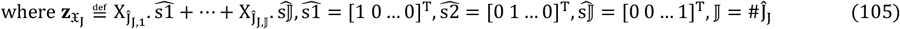

The algebraic model presented is J-reactant dependent and considers reactants as “siphons” where the contributions of the participating reactants are described as intermittent (**Table 1**) [28–30].

##### Constraint 16

(**Cs16**): The set of J-reactant dependent stoichiometry number matrices is an 𝕩-indexed (𝕩 = 1, …, 𝕏) 𝕏-membered 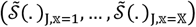 ordered set of constrained, unique and unequal stoichiometry number matrices [**From Cs15, T1, C1, C2**],

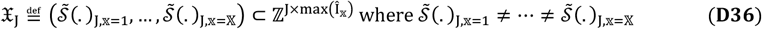

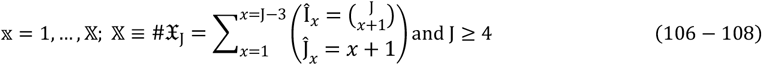

We simplify the notation without any loss of generality,

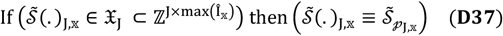

#### 2.2.3 Constraints to describe and incorporate indices of biochemical function

Whilst the Constraints described vide supra will ensure that our algebraic framework retains biochemical relevance, we will, nonetheless require direct indices of biochemical function by which we can model and characterize network-redundancy.

##### Constraint 17

**(Cs17)**: The null space for any 𝕩-stoichiometry number matrix 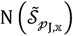 represents a biochemical network at equilibrium,

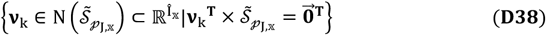

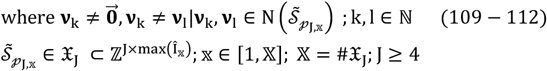

Biochemical networks can be “trivially”-perturbed as in sequestrations (complex formation, membrane-bound compartmentalization), post-translational modifications (acetylation, glycosylation, phosphorylation, etc.) and single nucleotide polymorphism (SNPs) variants of a reactant. Our rationale to preclude the “non-trivial” subset (deleterious mutations) is that these events are extremely rare and will alter the biochemical network irrevocably disallowing any meaningful comparison [7, 43].

##### Constraint 18

**(Cs18)**: The 𝕩^th^-stoichiometry number matrix 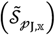 is perturbed 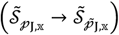 with *ρ*-PRVs and be studied by comparing the number and proportion of shared pairs of PRVs,

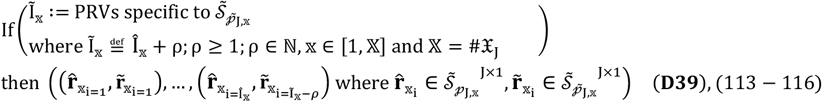

##### Constraint 19

(**Cs19**): There exists a bijection between the 𝕩^th^-stoichiometry number matrix (J-reactants, Î_𝕩_-PRVs) and the 𝕩^th^-biochemical network (J-reactants, Î_𝕩_-reactions),

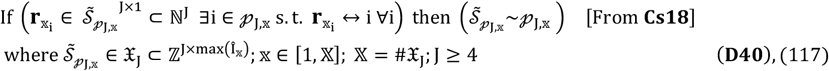

##### Constraint 20

**(Cs20)**: The probable dissociation constant for the i^th^-reaction of an 𝕩-specific biochemical network (η_i_(𝒫_J,𝕩_)) can be classified as Boolean,

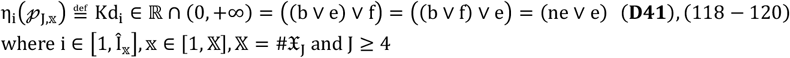

## 3 Methods

Constraints (**Cs1-Cs20**) and definitions (**D1-D41**), for a J-reactant dependent algebraic framework, will result in a set of 𝕩-indexed (𝕩 = 1, …, 𝕏) 𝕏-unique stoichiometry number matrices 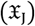 where each 𝕩-stoichiometry number matrix corresponds to an 𝕩-specific biochemical network (𝒫_J,𝕩_) (**Cs19**). Since we wish to model and characterize the redundant forms of a biochemical network, we seek 𝕩-indexed (𝕩 = 1, …, 𝕏) 𝕏-libraries of equivalent matrices where the 𝕩^th^-library can be mapped uniquely to the 𝕩^th^-stoichiometry number matrix and studied.

### 3.1 Strictly positive non-unitary rational number scalar multipliers as generators of a library of equivalent matrices for the 𝕩^th^-biochemical network

Stoichiometry coefficients are numerical representations of the reaction conditions and can be used to approximate the predicted/theoretical/assumed stoichiometry numbers for a set of reactants (**D1** and **D2** (**Eqs. (1–5)**). These can therefore, be fractions, negative numbers and zero, wherever applicable. Here, we use strictly positive ℚ_+_ (stoichiometry preserving) and non-unitary rational numbers as scalar multipliers for any 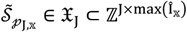 to generate an 𝕩-network specific library of equivalent matrices. Although the set of rational numbers (ℚ) is uncountable, we will restrict our consideration to a countable infinite subset with dimensions W × Y where W, Y ∈ ℕ (**Eqs. (121-155)**; **D42-D59**).

#### Theorem 3

(**T3**; **without proof**): The set of non-zero and strictly positive rational number multipliers is modelled as the square matrix,

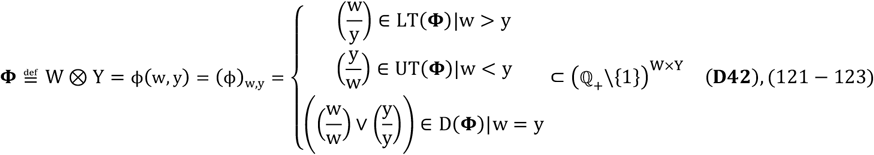

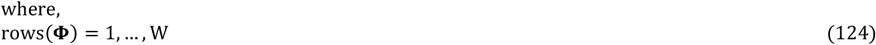

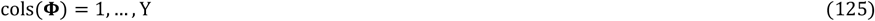

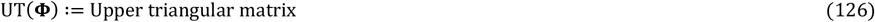

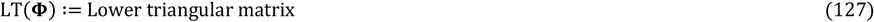

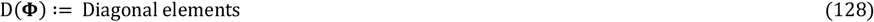

#### Corollary 3

(**C3; STx1: Eq. (8–20)**): The matrix of non-zero non-unitary rational number multipliers (**Ф** ⊂ (ℚ _+_\{1}) ^W ×Y^) is characterized by,

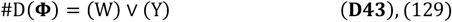

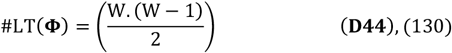

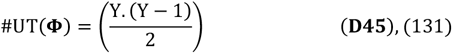

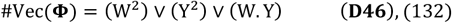

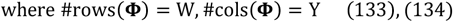

#### Corollaries 4-8

(**C4-C8; STx1: Eqs. (21–61)**): The following general results hold true for **Ф**,

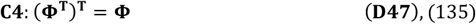

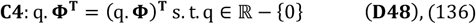

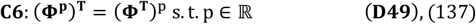

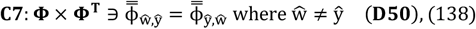

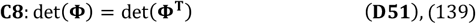

#### Lemma 1

(**L1**; **without proof**): The scalar product of a transposable pair of elements of **Ф** (off-diagonal pair) is unity,

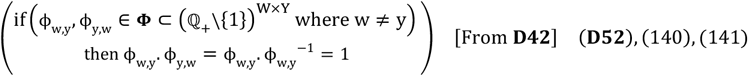

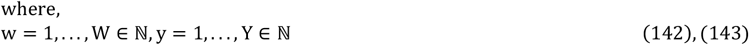

#### Theorem 4

(**T4; STx1: Eq. (62–71)**): The scalar product of all the off-diagonal elements of **Ф** is numerically equal to unity,

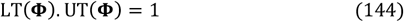

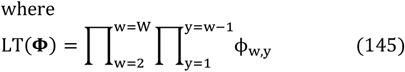

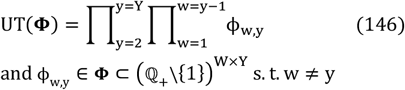

If we choose only off-diagonal elements of (ℚ_+_\{1}^W × Y^) as rational number multipliers for 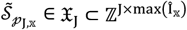, we can ensure the bijection with an 𝕩-biochemical network-specific *a*-indexed 𝔸-membered library of unique and equivalent rational number matrices 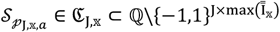. The rational number reaction vectors (RRVs; 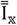) for 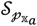 are then simply, transformed PRVs (Î_𝕩_) of 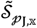,

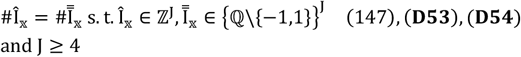

#### Theorem 5

(**T5**; **without proof**): A strictly positive and non-unitary rational number multiplier of 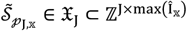 will generate an equivalent pair of rational number matrices,

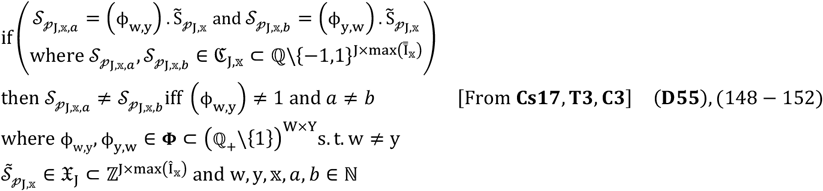

#### Theorem 6

(**T6; without proof**): Any 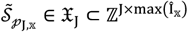 where 𝕩 ∈ [1, 𝕏] and 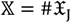 whence multiplied by strictly positive and non-unitary rational number multipliers will result in a library of equivalent rational number matrices for the 𝕩^th^**-**biochemical network,

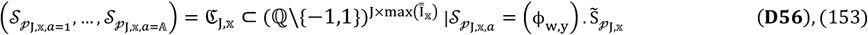

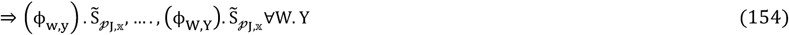

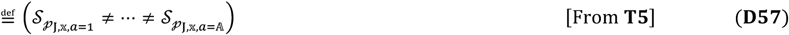

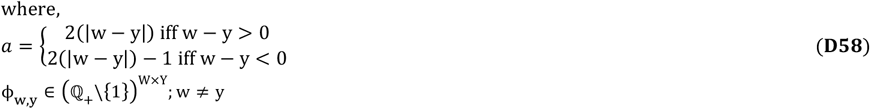

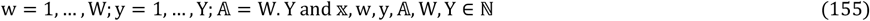

#### Corollary 9

(**C9**; **without proof**): For any 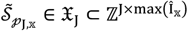 we have a J-reactant dependent 𝕩-library of *a*-indexed 𝔸-redundant forms for the 𝕩^th^-biochemical network,

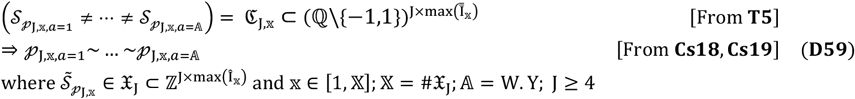

### 3.2 Probable dissociation constants, as parameters, to interrogate specific biochemical pathways

We utilize “ReDirection” to conduct detailed computational studies on known and well-characterized biochemical pathways [7]. “ReDirection” is an R-package that deploys a mathematically rigorous algorithm to compute the probable dissociation constant and assign an outcome to every reaction of a user-defined biochemical pathway [7, 8]. Briefly, “ReDirection” exhaustively checks a network-specific stoichiometry number matrix for linear dependency (rows, columns), lower bounds (reactants, PRVs) and reaction vector orientation (rows, columns) [7]. “ReDirection” then serially generates null space-generated subspaces by removing all redundant and trivial vectors, populating a reaction-specific sequence vector with numerical values drawn from each row and across all columns of this subspace and thence summing (‖. ‖_1_) for the same [7, 8]. These steps are done iteratively and recursively until the following conditions are met,

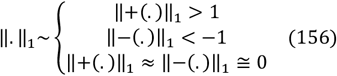

For each iteration thereafter, “ReDirection” will compute reaction-specific sequence vector descriptors such as the arithmetic mean, standard deviation, etc., screen each term and bin them to outcome-specific subsets whose terms are, in turn, summed and mapped to a positive real number [7]. These constitute the i^th^-reaction-specific output vector and its p1-norm is the probable dissociation constant (η_i_ ∈ ℝ ∩ (0, +∞)) [7, 8]. The aforementioned steps are continued until every reaction (∀i) has been assigned an unambiguous outcome in accordance with **Eq. (156)**.

### 3.3 Evaluating the consequences of introducing *ρ*-reactions into a specific biochemical pathway

The introduction of *ρ*-(reactions/RRVs) into an *a*-indexed redundant form of the 𝕩^th^-biochemical network 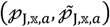 will be used as input to “ReDirection” since 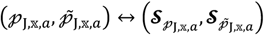 (**A19, C9**) (**Figure 2**). The probable dissociation constants will be computed and used to unambiguously assign outcomes to every reaction. In order to ascertain the effects of *ρ* ≥ 1-reactions, we will need to compute the number of shared reactions (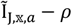; **A19, Figure 2**). From this subset, we compute the number, distribution and proportion of matched (m)-reactions which are non-equivalent (mne) and equivalent (me) (**Eqs. (157–160)) (Table 1; D60-D69**),

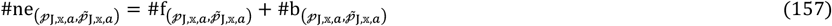

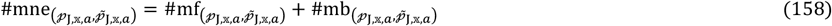

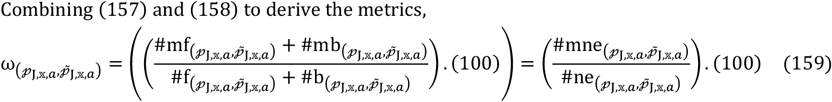

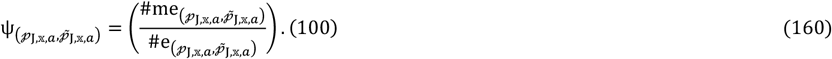

**Figure 1:**
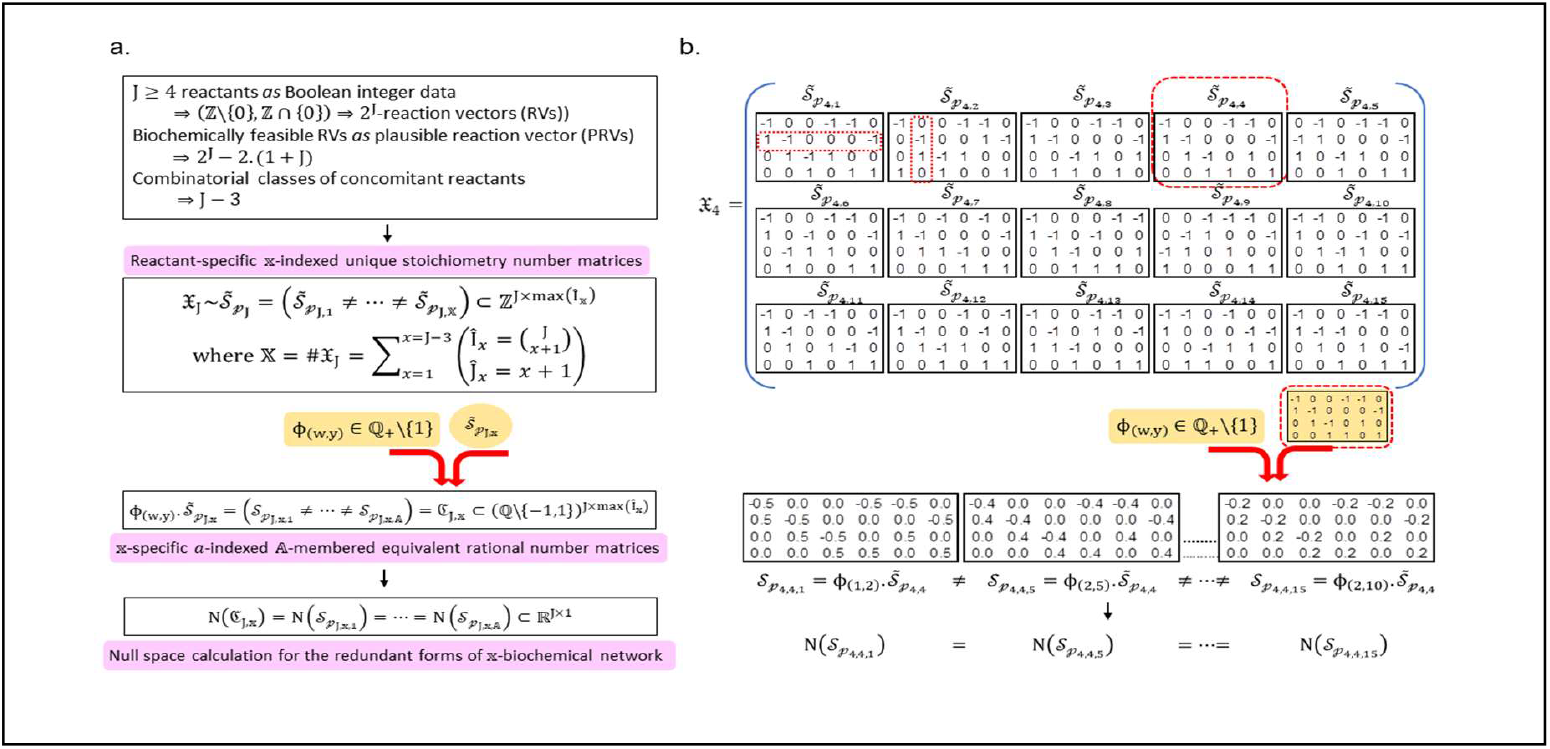
Construction and analysis of a reactant-dependent algebraic framework to model and characterize the redundant forms of a biochemical network: An algebraic framework to model and characterize redundant forms of a reactant-dependent biochemical network is constructed and populated in accordance with biochemically relevant paradigms. The framework is mathematical rigorous and comprises a set of unique stoichiometry number matrices. Each matrix is constrained by a lower bound for the number of reactants and thence the selected PRVs, is populated by integral changes which are Boolean (signed non-zero, zero) and combinatorially distributed across selected PRVs and represents a biochemical network. In order to study the redundant forms of a biochemical network, the dot product of a stoichiometry number matrix with non-zero non-unitary rational number multipliers is taken. The rational number matrices that result from these computations are similarly constrained, equivalent, degenerate with respect to a shared null space and comprise the redundant forms of a modelled biochemical network (CRaTER𝕩). The matrices have well defined algebraic properties and the pathways that they represent are easily parameterized in terms of probable dissociation constants and trivially perturbed. **Abbreviations**: **CRaTER**𝕩, Library of 𝕩-stoichiometry number matrix 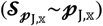 specific *a*-indexed 𝔸-membered rational number matrices 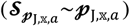 which are similarly constrained, equivalent, unequal and null-space redundant; 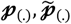, unperturbed- and perturbed-forms of a biochemical network; **PRV**, Plausible reaction vector;

**Figure 2:**
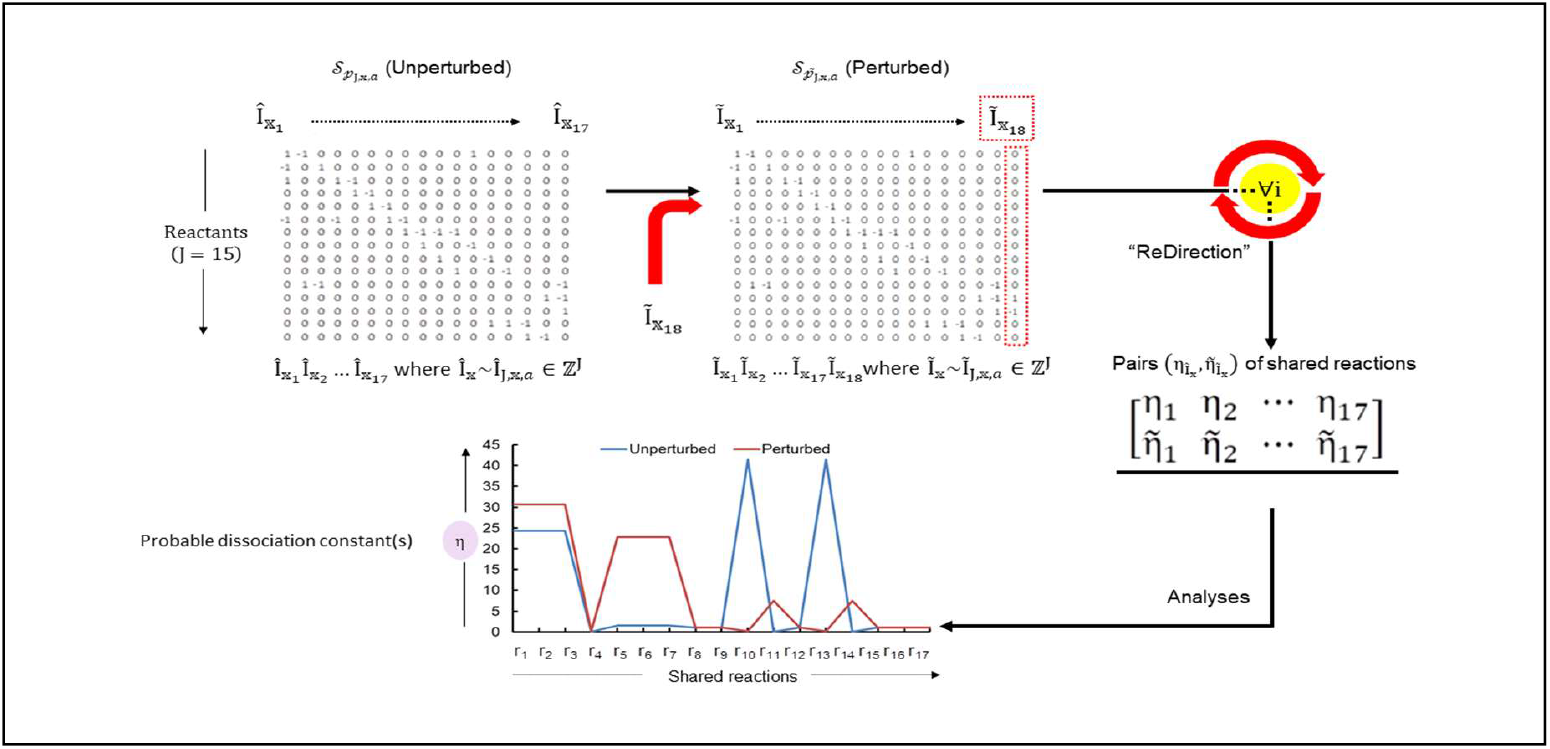
Perturbation studies for parameter selection and assessment of relevance of the redundant forms of a biochemical network: A perturbation is modelled by the inclusion of one or more extraneous reactions and its effect on the modelled biochemical network is studied by the pairwise comparison of the probable dissociation constants between shared reactions. This is possible since there is a one-one correspondence between a pathway (reactants, reactions) and its encoding rational number matrix (reactants, RRVs). The probable dissociation constant is a purely numerical measure that is computed by the R-package “ReDirection” for every reaction of a user-defined biochemical pathway. The metrics for these analyses are the numbers and proportions (matched, unmatched) of equivalent and non-equivalent reactions between the unperturbed- and perturbed-versions of a redundant form of an 𝕩^th^-biochemical network (*a*-indexed CRaTER𝕩-matrix). These network-derived characteristics are reformulated as metrics to compare the redundant forms of a biochemical network. These include 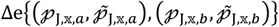 for pairs of concurrently considered redundant forms and 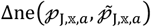 for single instances of the same. Both of these are computed by populating the appropriate confusion matrices with reaction-outcome data, after perturbing the system(s) and computing the indices (precision, accuracy, recall) which are then assessed. **Abbreviations**: **CRaTER**𝕩, Library of 𝕩-stoichiometry number matrix 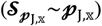 specific *a, b* indexed 𝔸-membered rational number matrices 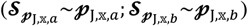 which are similarly constrained, equivalent, unequal (*a* ≠ *b*) and null-space redundant; 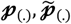, unperturbed- and perturbed-forms of a biochemical network; Δ**e**, Δ**ne**, Change in fraction of reactions with equivalent (e)**-** or non-equivalent (ne)**-**outcomes of a biochemical network after being perturbed; **PRV**, Plausible reaction vector; **RRV**, Rational number reaction vector;

### 3.4 Computational studies to evaluate the consequences of introducing *ρ*-reactions into known biochemical pathways

Biochemical pathways for the urea cycle and folate metabolism will be analysed with “ReDirection” [7, 8]. These are well characterized, conserved across taxa and make significant contributions to the intracellular milieu and thereby to the physiological function of a cell.

The human urea cycle is an important pathway in the utilization and removal of non-protein nitrogen (ammonia) and occurs, partly, in the mitochondria and the cytosol of the liver and kidneys. The inability to excrete urea over a prolonged period of time predisposes an individual to varying grades of uremia and encephalopathy. The clinical relevance of the urea cycle notwithstanding, Ornithine, a critical intermediate of the urea cycle is a significant contributor to polyamine biosynthesis (Putrescine, Spermidine, Spermine) [50, 51]. Polyamines (PAs) contribute non-trivially to recombination-mediated DNA repair, inwardly-rectifying potassium channels (K_ir_), cardiac conduction, cell proliferation and macrophage polarization [52–55]. The biochemical relevance of PAs is offset by their propensity to form free radicals *in vivo* and a non-trivial contribution to dysregulated cell growth [52–55]. Endogenous production of PAs is stringently regulated by the rate limiting enzymes Ornithine (EC 4.1.1.17)- and S-adenosylmethionine (EC 4.1.1.50)-decarboxylases [55, 56]. Here, we examine the effects of a multi-step and indirect reaction(s) (N-Acetylglutamate-, L-Glutamate-, 4-hydroxyglutamate-semialdehydes; 1-pyrroline-5- and 1-pyrroline-3-hydroxy-5-carboxylates) from Arginine, Proline and Glutamate (r_12_) on an integrated biochemical network of urea metabolism and polyamine biosynthesis. Folate metabolism connects disparate reactants (macromolecular, micronutrients) such as proteins, carbohydrates, lipids, DNA/RNA and vitamins. The “Folate trap” is a clinicopathological condition that is characterized by relative folate deficiency in the presence of vitamin B12 deficiency [57, 58]. Since folate is an important contributor to 1C-metabolism and DNA synthesis, its deficiency will result in macrocytic anemia (large and immature RBCs). In its role as a cofactor, folate is required by enzymes involved in the metabolism of several amino acids (Serine, Glycine, Histidine, Cysteine) [57, 58]. The direct conversion of dUMP to dTMP (r_18_) is likely to alter the flux of various reactants in an integrated biochemical network for folate and methionine metabolism and will be investigated in this study. Thymidine is required for DNA synthesis and the enzyme responsible for this conversion is thymidylate synthase (TS; EC 2.1.1.45) [58, 59]. Defects of this enzyme will result in the accumulation of dUMP and other deoxynucleotides which can damage DNA. Conversely, TS is a known contributor in the pathogenesis of hepatocellular carcinoma and development of chemoresistance [57–59].

## 4 Results

Here, we will investigate algebraic properties of the 𝕩^th^-biochemical network-specific library of equivalent rational number matrices, discuss the implications of introducing and thence perturbing known biochemical networks and evaluating parameter selection strategies to characterize network-level redundant forms of a biochemical network.

### 4.1 Defining and elucidating the properties of a library of equivalent rational number matrices for the 𝕩^th^-biochemical network

#### Corollaries-10

(**C10**) and -11 (**C11**) follow readily and is presented without formal proofs,

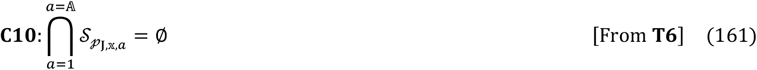

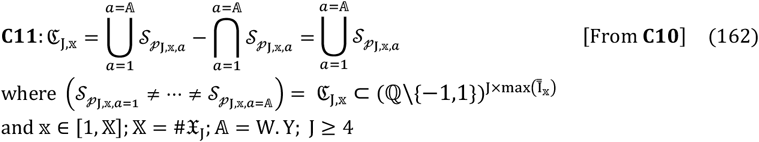

The null space is a numerical construct which is used to model a system of biochemical reactions at equilibrium (**A15**). The null space-generated subspace has also been shown to be biochemically relevant with the computation of the probable dissociation constant for every reaction of a user-defined biochemical network [7, 8].

#### Theorem 7

(**T7; STx1: Eqs. (72–84)**): The null spaces generated by members of ℭ_J,𝕩_ are equal,

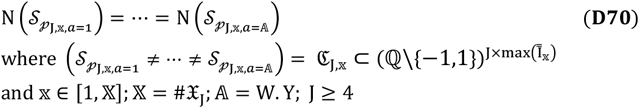

#### Theorem 8

(**T8; STx1: Eqs. (85-100)**): The linear map between ℭ_J,𝕩_ and the corresponding null space is many-to-one,

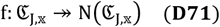

In accordance with **T7** and **T8** we refine our definition of ℭ_J,𝕩_ to include this degeneracy (D) (**Figure 1**). In other words,

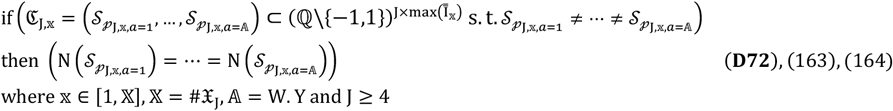

We will now present a refined definition for an *a*-indexed 𝔸-membered library of constrained, rational number, equivalent and redundant matrices for the 𝕩^th^-biochemical network (CRaTER𝕩),

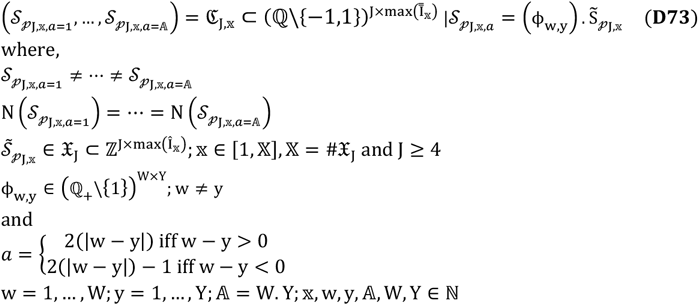

Generic mathematical properties for ℭ_J,𝕩_ include being rectangular (4 ≤ J < max(Î_𝕩_); **Eq**. (**165**)) and comprising reaction vectors that are linearly dependent.

#### Lemma 2

(**L2; without proof**): The set 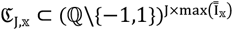 is commutative or Abelian with regards to addition,

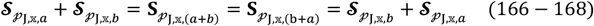

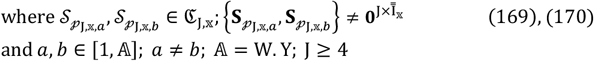

#### Theorem 9

(**T9; STx1: Eqs. (101–108)**): The set 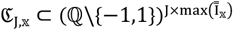 is a semigroup with respect to addition and is conditionally Abelian,

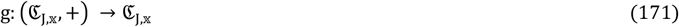

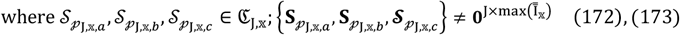

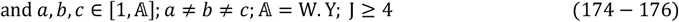

### 4.2 “ReDirection”-based analysis of known biochemical pathways

We now complement our theoretical assertions with detailed computational studies on pathways for urea synthesis and folate metabolism. Perturbations are introduced as single extraneous reactions and the probable dissociation constants for each pair of shared reactions (unperturbed, perturbed) is computed 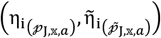 [7, 8].

The data, for the urea cycle and thence polyamine biosynthesis suggests that there is a decrease in the absolute number of equivalent reactions (**Figures 3a and 3b, Table 3**). The increase in Glutamate synthesis from Arginine 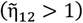 will stimulate the synthesis of Ornithine 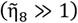 which will condense with urea and form Arginine 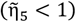 (**Figures 3a and 3b, Table 2)**. This, in turn, will drive the urea cycle via the formation of Citrulline, Aspartate and Fumarate 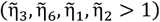 (**Figures 3a and 3b, Table 3)**. These modifications are compounded by reductions in the probable dissociation constants for the syntheses of Glutamate from Fumarate and thence Aspartate 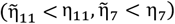 (**Figures 3a and 3b, Table 3)**. Our results also demonstrate a heightened synthesis of Ornithine via Glutamate 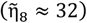. This, we combined with the futile utilization via Citrulline 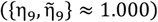 and synthesis to Arginine 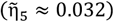 directly will ensure increased availability of Ornithine and thence PAs (**Figures 3a and 3b, Table 3)**.

**Table 2:** Numerical case studies for a J-reactant dependent algebraic model to model and characterize the redundant forms of a biochemical network.

| Reactants<br>(J) | RV<br>(2<sup>J</sup>) | PRVs<br>(2<sup>J</sup> - 2(1 + J)) | Classes<br>(J - 3 (J)) | Set(s) of reactants<br>($\bar{\theta}_{J_j}$) | Set(s) of PRVs<br>($\hat{I}_{J_j}$) | Set(s) of SNMs<br>($\# \mathfrak{X}_J = \mathbb{X}$) | CraTER<sub>x</sub><br>($\# \mathfrak{C}_{J,x} = \mathbb{A}$)<br>($x \in [1, \mathbb{X}]$) |
| --- | --- | --- | --- | --- | --- | --- | --- |
| 4 | 16 | 6 | 1 | $\bar{\theta}_{J_4} = \{\bar{\theta}_{J_{4,1}} = 2\}$ | $\hat{I}_{J_4} = (\hat{I}_{J_{4,1}} = 6)$ | $\# \mathfrak{X}_4 = \left( \mathfrak{S}_{J_{4,1}}(2,6): \binom{6}{2} = 15 \right)$ | $\mathfrak{C}_{J,x,a}$<br>$a = 1, \dots, \mathbb{A}$<br>$x \in [1, 15]$ |
| 5 | 32 | 20 | 2 | $\bar{\theta}_{J_5} = \begin{pmatrix} \bar{\theta}_{J_{5,1}} = 2 \\ \bar{\theta}_{J_{5,2}} = 3 \end{pmatrix}$ | $\hat{I}_{J_5} = \begin{pmatrix} \hat{I}_{J_{5,1}} = 10 \\ \hat{I}_{J_{5,2}} = 10 \end{pmatrix}$ | $\# \mathfrak{X}_5 = \left( \mathfrak{S}_{J_{5,1}}(2,10): \binom{10}{2} = 45 \right. \\ \left. \mathfrak{S}_{J_{5,2}}(3,10): \binom{10}{3} = 120 \right) = 165$ | $\mathfrak{C}_{J,x,a}$<br>$a = 1, \dots, \mathbb{A}$<br>$x \in [1, 165]$ |
| 6 | 64 | 50 | 3 | $\bar{\theta}_{J_6} = \begin{pmatrix} \bar{\theta}_{J_{6,1}} = 2 \\ \bar{\theta}_{J_{6,2}} = 3 \\ \bar{\theta}_{J_{6,3}} = 4 \end{pmatrix}$ | $\hat{I}_{J_6} = \begin{pmatrix} \hat{I}_{J_{6,1}} = 15 \\ \hat{I}_{J_{6,2}} = 20 \\ \hat{I}_{J_{6,3}} = 15 \end{pmatrix}$ | $\# \mathfrak{X}_6 = \left( \mathfrak{S}_{J_{6,1}}(2,15): \binom{15}{2} = 105 \right. \\ \mathfrak{S}_{J_{6,2}}(3,20): \binom{20}{3} = 180 \\ \left. \mathfrak{S}_{J_{6,3}}(4,15): \binom{15}{4} = 1365 \right) = 1650$ | $\mathfrak{C}_{J,x,a}$<br>$a = 1, \dots, \mathbb{A}$<br>$x \in [1, 1650]$ |
| 7 | 128 | 112 | 4 | $\bar{\theta}_{J_7} = \begin{pmatrix} \bar{\theta}_{J_{7,1}} = 2 \\ \bar{\theta}_{J_{7,2}} = 3 \\ \bar{\theta}_{J_{7,3}} = 4 \\ \bar{\theta}_{J_{7,4}} = 5 \end{pmatrix}$ | $\hat{I}_{J_7} = \begin{pmatrix} \hat{I}_{J_{7,1}} = 21 \\ \hat{I}_{J_{7,2}} = 35 \\ \hat{I}_{J_{7,3}} = 35 \\ \hat{I}_{J_{7,4}} = 21 \end{pmatrix}$ | $\# \mathfrak{X}_7 = \left( \mathfrak{S}_{J_{7,1}}(2,21): \binom{21}{2} = 210 \right. \\ \mathfrak{S}_{J_{7,2}}(3,35): \binom{35}{3} = 6545 \\ \mathfrak{S}_{J_{7,3}}(4,35): \binom{35}{4} = 52,360 \\ \left. \mathfrak{S}_{J_{7,4}}(5,21): \binom{21}{5} = 20,349 \right) = 79464$ | $\mathfrak{C}_{J,x,a}$<br>$a = 1, \dots, \mathbb{A}$<br>$x \in [1, 79464]$ |

**Abbreviations:**
J: Reactants RV: Reaction vector PRV: Plausible reaction vector SNM: Stoichiometry number matrix CraTER<sub>x</sub>: Constrained, rational number, equivalent and redundant matrices for the $x^{\text{th}}$ -biochemical network

**Table 3:**
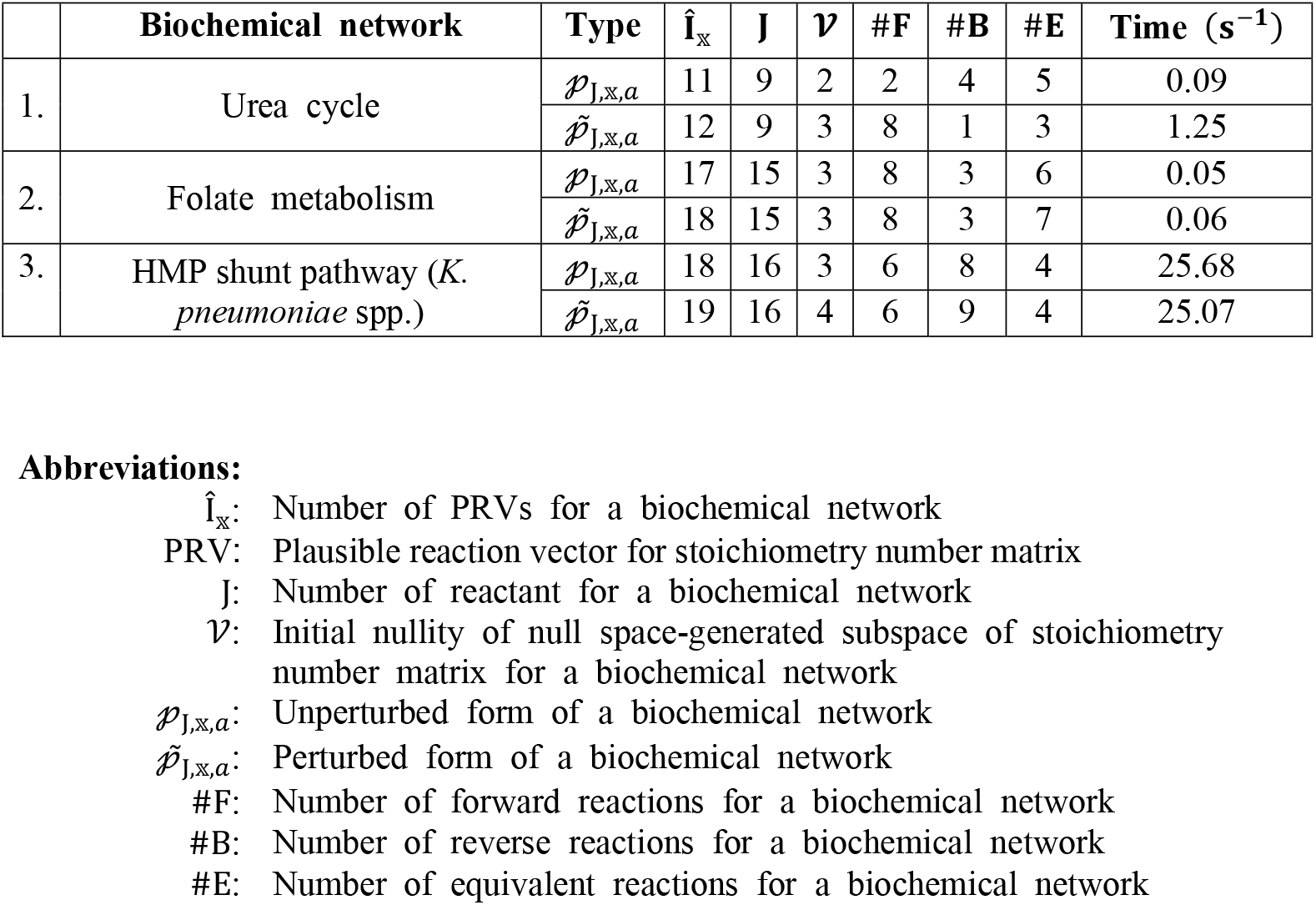
“ReDirection”-based analysis for a redundant form (unperturbed, perturbed) of known biochemical networks.

| | Biochemical network | Type | $\hat{\mathbf{I}}_{\mathbf{x}}$ | $\mathbf{J}$ | $\mathcal{V}$ | #F | #B | #E | Time ($\text{s}^{-1}$) |
| --- | --- | --- | --- | --- | --- | --- | --- | --- | --- |
| 1. | Urea cycle | $\mathcal{P}_{\mathbf{J},\mathbf{x},a}$ | 11 | 9 | 2 | 2 | 4 | 5 | 0.09 |
| | | $\tilde{\mathcal{P}}_{\mathbf{J},\mathbf{x},a}$ | 12 | 9 | 3 | 8 | 1 | 3 | 1.25 |
| 2. | Folate metabolism | $\mathcal{P}_{\mathbf{J},\mathbf{x},a}$ | 17 | 15 | 3 | 8 | 3 | 6 | 0.05 |
| | | $\tilde{\mathcal{P}}_{\mathbf{J},\mathbf{x},a}$ | 18 | 15 | 3 | 8 | 3 | 7 | 0.06 |
| 3. | HMP shunt pathway (*K. pneumoniae* spp.) | $\mathcal{P}_{\mathbf{J},\mathbf{x},a}$ | 18 | 16 | 3 | 6 | 8 | 4 | 25.68 |
| | | $\tilde{\mathcal{P}}_{\mathbf{J},\mathbf{x},a}$ | 19 | 16 | 4 | 6 | 9 | 4 | 25.07 |

**Abbreviations:**
- $\hat{\mathbf{I}}_{\mathbf{x}}$ : Number of PRVs for a biochemical network - PRV: Plausible reaction vector for stoichiometry number matrix - $\mathbf{J}$ : Number of reactant for a biochemical network - $\mathcal{V}$ : Initial nullity of null space-generated subspace of stoichiometry number matrix for a biochemical network - $\mathcal{P}_{\mathbf{J},\mathbf{x},a}$ : Unperturbed form of a biochemical network - $\tilde{\mathcal{P}}_{\mathbf{J},\mathbf{x},a}$ : Perturbed form of a biochemical network - #F: Number of forward reactions for a biochemical network - #B: Number of reverse reactions for a biochemical network - #E: Number of equivalent reactions for a biochemical network

**Figure 3:**
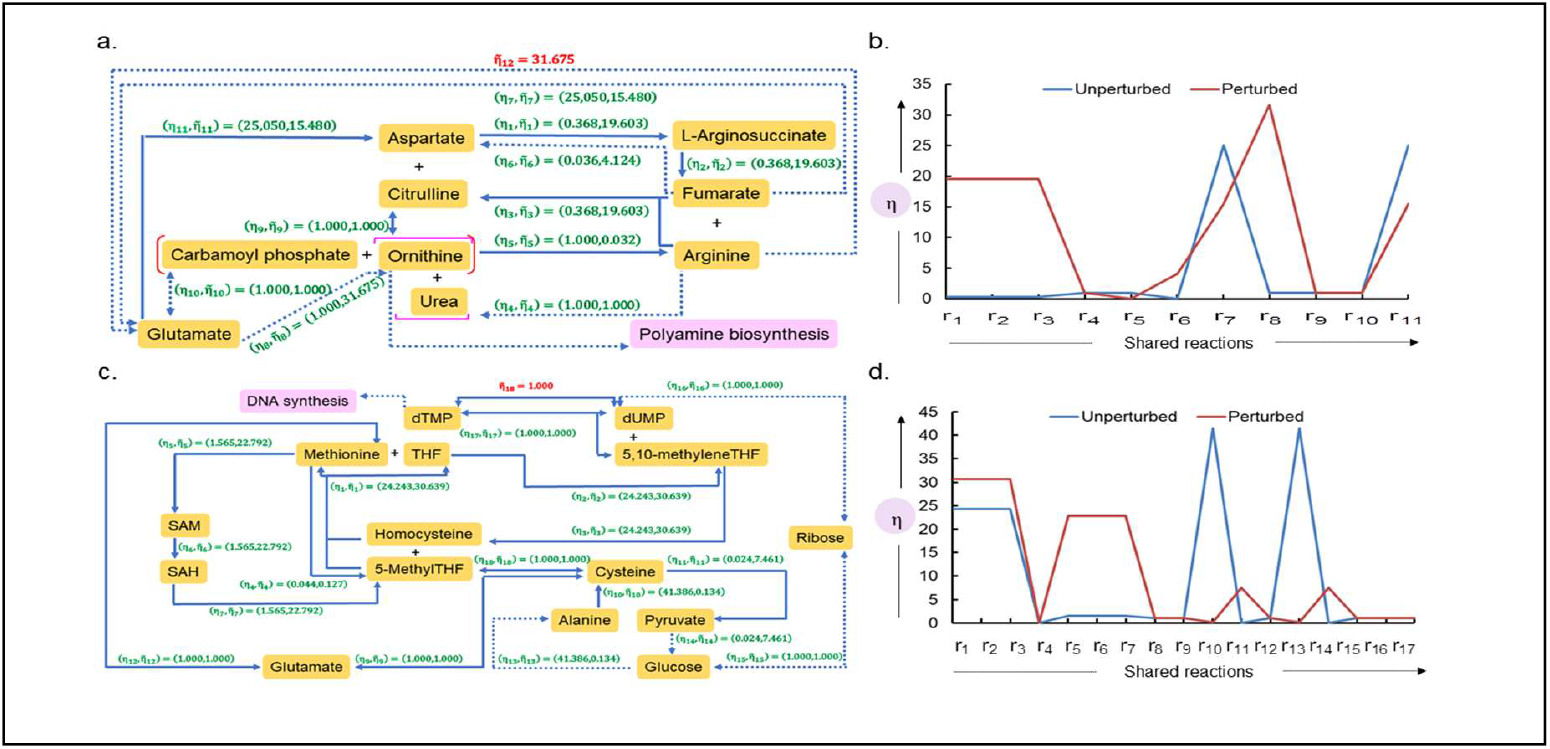
Computational studies to model, evaluate and establish biochemical and physiological relevance for a reactant-dependent algebraic framework: **a), b)** The urea cycle is characterized by several shunt pathways via Aspartate (Kreb’s cycle, r_6_) and Carbamoyl phosphate/Glutamate (Kreb’s cycle, r_7_; D-amino acids, r_5_; Arginine-Proline, r_8_). The synthesis of Citrulline (r_3_) with Nitric oxide synthase (NOS, EC1.14.13.39) utilizes NADPH and nitric oxide which is an important member of reactive oxygen (RO)- and reactive nitrogen (RN)-species (ROS, RNS). In this study, we investigate the effects of converting Arginine to Glutamate (r_12_) and thence to urea and Ornithine. Ornithine, in turn, is a critical precursor in the biosynthetic pathway of the polyamines (Putrescine, Spermidine, Spermine). These compounds are involved in DNA synthesis, cell proliferation and growth and the regulation of a subclass of potassium receptors and **c), d)** The folate cycle (THF, 5-methylTHF and 5,10-methyleneTHF) is responsible for the movement and exchange of several small molecules (methyl groups, vitamins, amino acids) between DNA/RNA, proteins and lipids. We examine effects of a direct and equally probable interconversion between dTMP to dUMP (r_18_) on a biochemical network of folate metabolism. **Abbreviations**: **CRaTER**𝕩, Library of 𝕩-stoichiometry number matrix 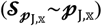 specific *a*-indexed 𝔸-membered rational number matrices 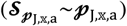 which are similarly constrained, equivalent and redundant; 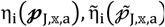, probable dissociation constant for a shared reaction between pathways or redundant forms of a biochemical networks; **mne**, shared and matched non-equivalent reactions; **me**, shared and matched equivalent reactions; **THF**, Tetrahydrofolate;

We also observe and study the relevance of interconversion dUMP and dTMP 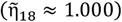, in folate metabolism, is equivalent and likely to be a key regulatory step. We also note that although the proportion of equivalent and non-equivalent reactions remains unchanged, the magnitudes of the probable dissociation constants differ significantly (**Figures 3c and 3d, Table 3**). Amongst the reactions that exhibit a complete reversal are the reactants Cysteine, Alanine, Pyruvate and Glucose 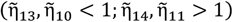 (**Figures 3c and 3d, Table 3**). On the other hand, alterations in the magnitude of the probable dissociation constants are also observed for reactions that involve methionine, S-adenosylmethionine, Homocysteine, S-adenosylhomocysteine and the tetrahydrofolates 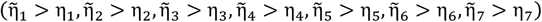 (**Figures 3c and 3d, Table 3**). Since 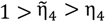, a greater quantity of Homocysteine may be available for its reaction with 5-methylenetetrahydrofolate 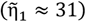 as opposed to the simple reversal to Methionine 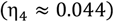. The ensuing reactions converts tetrahydrofolate to 5,10-methylenetetrahydrofolate 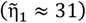 and thence to dTMP via methylation of dUMP. A critical mediator favouring this step is the availability of Ribose via Glucose, Alanine and Pyruvate (Glucose/Alanine/Cysteine, Cysteine/Glucose/Pyruvate) (**Figures 3c and 3d, Table 3**). Since 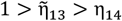, the conversion of Glucose to Alanine 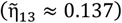 is much slower as compared to the same via Pyruvate (η_14_ ≈ 0.024). This results a greater availability of Glucose for Ribose synthesis and subsequent ribosylation of the de novo synthesized Thymine base (Thymidine). This reactant is critical in the synthesis of the Uridine-Thymidine transversion and thence DNA synthesis and a perturbation will shift the equilibrium towards compensatory synthesis (**Figures 3c and 3d, Table 3; ST2, SD2**).

### 4.3 Perturbation-based metrics to characterize the redundant forms of a biochemical network

Although the *a*^th^-redundant form of an 𝕩^th^-biochemical network is easily evaluated (unperturbed, perturbed) 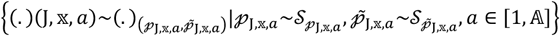 (**Eq. (177)**), we will need a metric that compares 𝔸-redundant forms of a biochemical network concurrently. Here, we will discuss the derivation of metric which be used to compare any pair of redundant forms 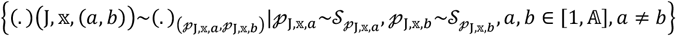 (**Eq. (178)**) (**Cs19, C9**). In order to accomplish this, we will reformulate previously derived metrics into non-local (less sensitive, numerically robust, specific) and local (greater sensitivity, reduced specificity, robustness)-versions (**D74, D75**).

Let us compute confusion matrices for the final outcomes between pairs of shared reactions of the unperturbed- and perturbed-forms of a biochemical network, which we will evaluate as parameters 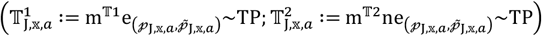 (**Eqs. (179–182)) (Table 1; D76-D83**),

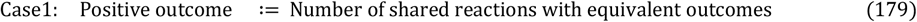

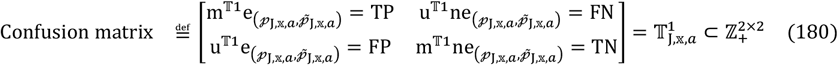

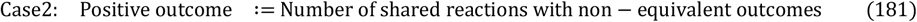

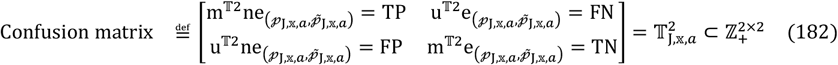

The indices of accuracy (Ac), precision (Pr), recall (Re), false positive rate (FPr) and specificity (SPy) will be computed (**Eqs. (183–190)**) (**Table 4**),

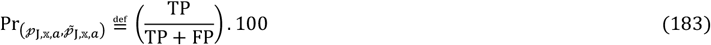

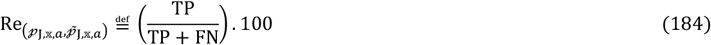

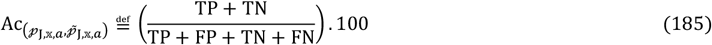

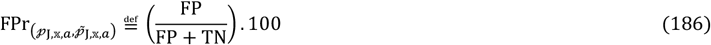

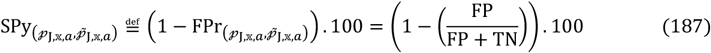

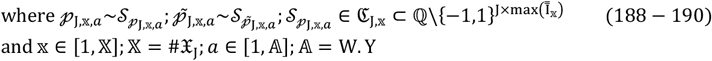

**Table 4:** Evaluating the confusion matrices for a redundant form (unperturbed, perturbed) of known biochemical networks.

| Confusion matrix | Network | TP | FP | FN | TN | Pr | Re | Ac | SPy |
| --- | --- | --- | --- | --- | --- | --- | --- | --- | --- |
| $\mathbb{T}_{j,x,a}^1$ | Urea | 3 | 2 | 4 | 2 | 60% | $\approx 43\%$ | $\approx 46\%$ | 50% |
| | Folate | 6 | 0 | 4 | 7 | 100% | 60% | $\approx 77\%$ | 100% |
| | HMP (K.pa<sup>+</sup>) | 4 | 0 | 5 | 9 | 100% | $\approx 44\%$ | $\approx 72\%$ | 100% |
| $\mathbb{T}_{j,x,a}^2$ | Urea | 2 | 4 | 2 | 3 | $\approx 33\%$ | 50% | $\approx 46\%$ | $\approx 43\%$ |
| | Folate | 7 | 4 | 0 | 6 | $\approx 64\%$ | 100% | $\approx 77\%$ | 60% |
| | HMP (K.pa<sup>++</sup>) | 9 | 5 | 0 | 4 | $\approx 64\%$ | 100% | $\approx 72\%$ | $\approx 44\%$ |

We can rewrite 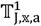 and 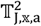 (**Eqs. (180), (182)**) in terms of the previously derived network-specific (**Eqs. (183–190**)) and generalize this as the bijection (g),

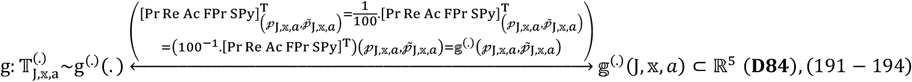

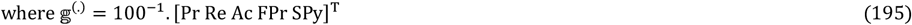

Let us now partition into non-local (𝕘^1^(.) ∈ 𝒢(ℭ_J,𝕩_))- and local (𝕘^2^(.) ∈ ℋ (ℭ_J,𝕩_))-parameters to evaluate and characterize the redundant forms of a biochemical network after a perturbation. Whilst the former will evaluate the shift or lack thereof, for reactions with equivalent outcomes between pairs of concurrently occurring redundant forms, the latter will characterize the alteration in flux of reactants/products through reactions with non-equivalent outcomes (forward, reverse) for any single redundant form. Paraphrasing the aforementioned assertions we can define,

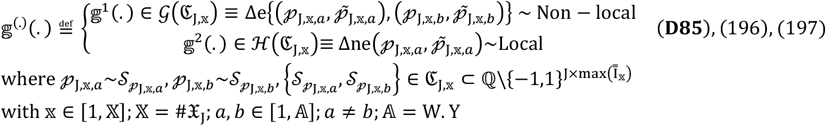

Expanding each of these and rewriting **Eqs**.-(**198**) and -(**199**) in accordance with **D84**,

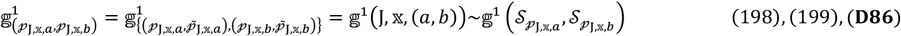

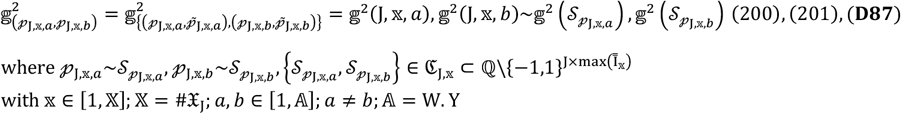

## 5 Discussion

The intracellular milieu is complex and any model must incorporate contributors to this complexity. Our algebraic framework can be used to model and characterize the redundant forms of a biochemical network with parameters that are both, theoretically sound and biochemically relevant.

### 5.1 Parameter selection to analyse the redundant-forms of a biochemical network

Although the outcome of reaction equivalence suggests data ambiguity for a reaction, inferring a regulatory role for the same is equally plausible. Conversely, reactions with non-equivalent (forward, reverse) outcomes are likely to represent the prevailing physicochemical conditions and may represent the response elements of a biochemical network. These hypotheses were examined, numerically, *post hoc* by the evaluation of 𝕘^(.)^(.) for the unperturbed- and perturbed-forms of the urea cycle and folate metabolism pathways (**D85**; **Table 4**).

The recall of a metric is a measure of how sensitive a pathway is to perturbation. Since these are almost identical (*Re*_(*Urea,Folate*)_ ≈ 52%) and of moderately low magnitude, the utilization of reaction equivalence as the relevant non-local parameter is easily justified (*SPy*_(*Urea,Folate*)_ ≈ 75%, *Ac*_(*Urea,Folate*)_ ≈ 62%) (**Table 4**). The precision or positive predictive value of a metric is an indicator of robustness upon change. This, too, for an outcome of reaction equivalence is a useful parameter to comprehend and compare biochemical pathways pre- and post-perturbation (*Pr*_(*Urea,Folate*)_ ≈ 80%) (**Table 4**). Interestingly, if the original confusion matrix was redefined with a reaction outcome of non-equivalence as *TP* and reaction equivalence as 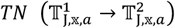 we get different results (**Table 4**). Since reactions with non-equivalent outcomes are likely to be influenced by the local intracellular milieu, this measure will be a poor choice whence comparing biochemical networks (*Re*_(*Urea,Folate*)_ ≈ 75%) (**Table 4**). However, since forward- and reverse-reactions regulate the flux of the reactants/products through a biochemical network, outcomes of non-equivalence for the majority of the reactions will result in a net direction for a pathway (*SPy*_(*Urea.Folate*)_ ≈ 52%) rather than robustness (*Pr*_(*Urea,Folate*)_ ≈ 49%) (**Table 4**).

We conclude, from these studies, that reactions with an equivalent outcome may constitute a useful metric of comparison between the redundant forms of a biochemical network whence taken concurrently. On the other hand, since the flux of reactants/products through reactions with a non-equivalent outcome (forward, reverse) is dependent on the prevailing physicochemical conditions, this metric is better suited to characterizing an instance of a redundant form of a biochemical network. Extending these results to 𝔸-redundant forms of an J-reactant dependent 𝕏^th^-biochemical network (CRaTER𝕏),

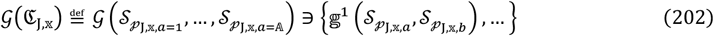

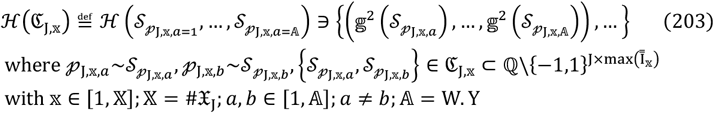

### 5.2 Biomedical relevance of the redundant forms of a biochemical network for the HMP-shunt pathway in the genesis of Klebsiella spp.-mediated formaldehyde resistance

Nosocomial (healthcare-associated) infections are usually observed in patients, in-residence, at a healthcare facility for a significant period of time [61]. Formaldehyde (HCHO) is a surface-disinfectant, a fixative and is also used to sterilize instruments such as endoscopes which cannot be autoclaved or heated. HCHO is also generated intracellularly as part of the respiratory burst (macrophages, neutrophils) in response to injury/infection, enzyme demethylation and tetrahydrofolate degradation [62–64]. The electrophilic reactivity of HCHO will result in adduct-formation and crosslinking (DNA/RNA/proteins) which is toxic to cells and non-methylotrophic bacteria [62, 63]. Interestingly, several studies have demonstrated that the absence of HCHO-assimilating genes does not preclude non-methylotrophic bacteria from utilizing and detoxifying the same [62–65]. In fact, gram negative bacteria possess several molecular mechanisms to sense, detoxify and thrive in the presence of HCHO (*NmlR*-*adhC*-*estD, FrmR*-*frmAB*) [62–65]. For example, a comparative analysis of several *Neisseria spp*. suggests that the greater virulence of *N. meningitidis* could be its rapid dissemination. This, in turn, could be on account of possessing both, *adhC* (alcohol dehydrogenase, EC1.1.2.84) and *estD* thioesterase, EC3.1.2.12) genes as opposed to *N. gonorrhoea* (*estD*)..[62, 63]. Similar mechanisms are operative for *Bacillus* spp. (*HxlR*-*hxlAB*) and *Klebsiella pneumoniae, Pseudomonas aeruginosa, Escherichia coli* (*FrmR*-*frmAB*) [63]. The biomedical and clinical relevance for these horizontally acquired genes was appreciated when it was discovered that certain multi-drug resistant (MDR) strains of *Klebsiella pneumoniae* (sequence type 4523; ST4523) and *Pseudomonas aeruginosa* were found to possess the FrmRAB operon and could thrive in an environment of HCHO along with other biocides [66, 67].

The molecular mechanisms to activate and thence assimilate HCHO depends on it condensing with reduced Glutathione (GSH) which in turn, is oxidized (GSSG) and will need to be reduced. The HMP-shunt pathway (oxidative, non-oxidative) is a conserved (eukaryotes, bacteria) pathway that produces reduced Nicotinamide adenine dinucleotide phosphate (NADPH + H^+^), phosphoribosyl pyrophosphate (PRPP) and many other potentially anaplerotic intermediates for Kreb’s cycle [68–71]. Although NADPH + H^+^ partakes in several important pathways, it is the major source of reducing equivalents in the reduction and subsequent regeneration of GSH [62–65]. Clearly the integrity and functionality of the HMP-shunt pathway is important for the growth and continual survival of bacteria, a factor facilitated by the concomitant formation of PRPP. The Ribulose-5-phosphate (RuMP)-pathway is either the bona fide oxidative phase of the pathway, as in methylotrophic bacteria (*M. aminofaciens, B. bravis, Bacillus* spp.) and some archaea (methanogens, methylotrophic), or complements an existent HMP-shunt pathway (non-methylotrophic organisms; *S. cerevisae, E. coli*) [64–67, 72, 73]. RuMP/XuMP condenses with formaldehyde to give a hexulose-6-phosphate which isomerizes into fructose-6-phosphate via the reversible reactions that are catalysed by hexulose-6-phosphate synthase (EC4.1.2.43) and phosphohexose isomerase (EC5.3.1.9) respectively [64–67].

Horizontal gene transmission (HGT) may contribute to HCHO-mediated resistance in nosocomial infections via the HMP-pathway. Although, the inter-conversion between β-D fructose 6-phosphate and D-ribulose 6-phosphate (r_19_) on a redundant form of the HMP-shunt pathway, in a multi-drug variant of *K. pneumoniae* (ST4523), does not alter the outcome of the shared equivalent reactions 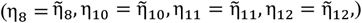, the same does not hold true for the shared non-equivalent reactions (**Figures 4a and 4b, Table 2; ST3, SD3**). These exhibit a reduction in magnitude 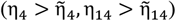 as well as reversed outcomes 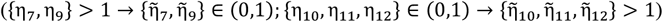 (**Figures 4a and 4b, Table 2; ST3, SD3**). From this data it is clear that any existing and host-specific Ribulose-5-phosphate is utilized (r_7,9_) and/or redirected (r_10–12_). This apparent loss of D-Ribulose-5-phosphate will, by the law of mass action and in concert with r_4_ and r_14_, result in additional D-Ribulose-5-phosphate being formed from D-Fructose-6-phosphate via 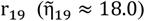 (**Figures 4a and 4b, Tables 3** and **4; ST3, SD3**). The subsequent condensation of HCHO by D-Ribulose-5-phosphate and thence assimilation/utilization into D-Fructose-6-phosphate will allow *K. pneumoniae* (ST4523) to grow, thrive and disseminate (**Figure 4c**).

**Figure 4:**
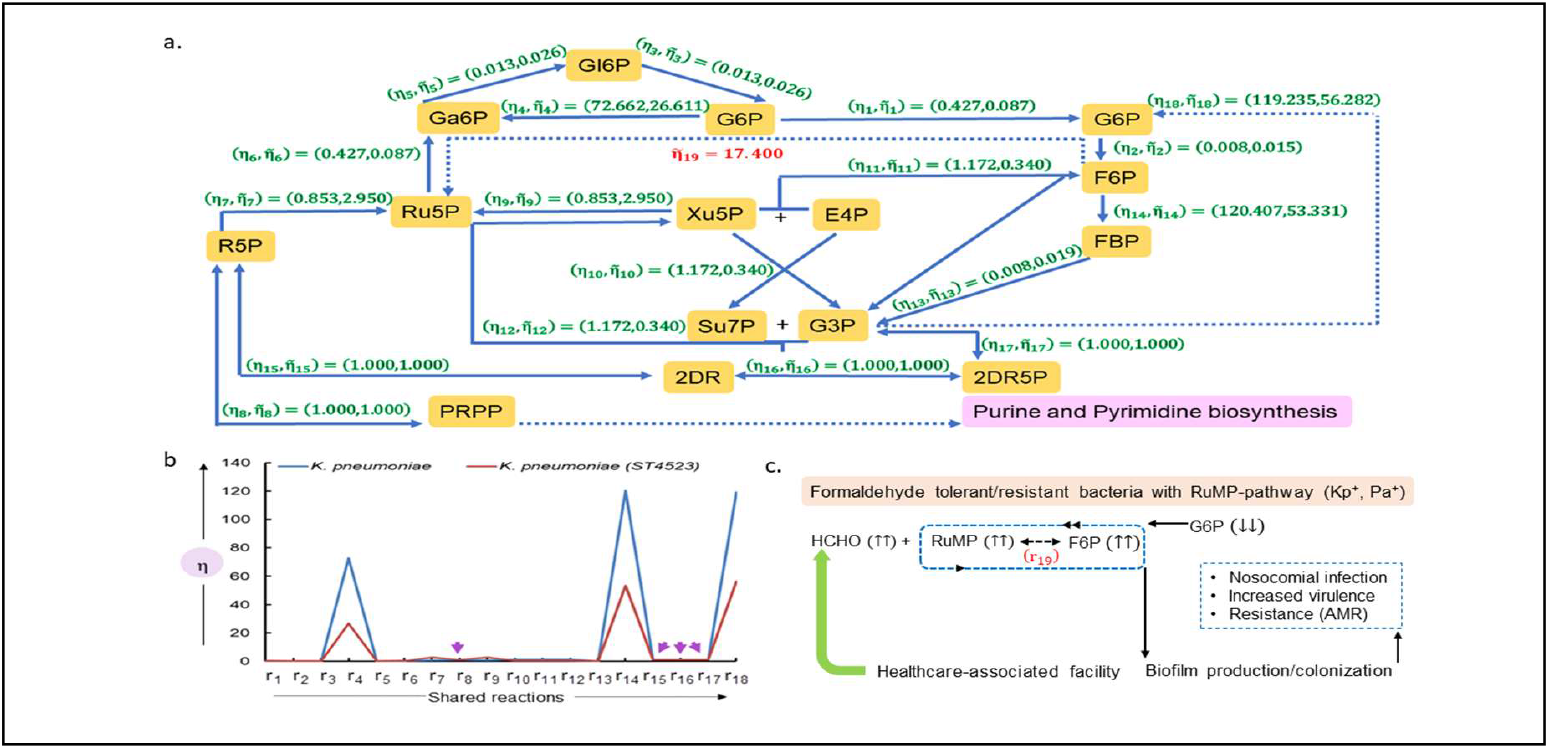
Biomedical relevance of the redundant forms of a biochemical network for the HMP-shunt pathway in the genesis of formaldehyde resistance: **a), b)** Analysis of a biochemical network of the hexose monophosphate shunt pathway in *K. pneumoniae* (ST4523) before and after the introduction of r_19_. In this study the direct conversion of β-D fructose 6-phosphate to D-ribulose-5-phosphate (r_19_) is investigated in terms of the magnitude and distribution of probable dissociation constants for the shared equivalent- and non-equivalent reactions of redundant forms of the biochemical network for the HMP-pathway (*Klebsiella pneumoniae, Klebsiella pneumoniae* ST4523). Although this conversion is absent in humans, the reaction is a significant contributor to formaldehyde fixation in bacteria and some archaea. The reactions occur via D-arabino hex-3-ulose 6-phosphate and includes the enzymes (6-phospho-3-hexuloisomerase, EC5.3.1.27; 3-hexulose-6-phosphate synthase, EC4.1.2.43) as independent polypeptides (archaea) or bifunctional protein (methylotrophic bacteria) and **c)** Schematic diagram for a model of enhanced nosocomial infection. Here, an increased virulence and resistance in *Klebsiella pneumoniae* and other non-methylotrophic bacteria such as *Pseudomonas aeruginosa* and *Escherichia coli* may occur if genes to support the RuMP pathway were horizontally acquired. These bacteria would then possess the RuMP-pathway in addition to the conserved HMP-shunt pathway and formaldehyde detoxifying genes (*adhC, estD*). This may result in bacteria that could thrive on formaldehyde exposure in a hospital setting (host-pathogen interface, formaldehyde-sterilized endoscopic instruments). **Abbreviations**: **EC**, enzyme commission; **F6P**, Fructose-6-phosphate; **G6P**, Glucose-6-phosphate; **NADPH**, reduced Nicotinamide adenine dinucleotide phosphate; **mM**, Millimolar; ( **Pa**^**+**^, **Kp**^**+**^), *Pseudomonas aeruginosa* and *Klebsiella pneumoniae* with horizontally acquired genes to support the RuMP pathway; **PRPP**, phosphoribosyl pyrophosphate; **RuMP**, Ribulose-5-phosphate;

## 6 Concluding remarks

This study develops a reactant-dependent algebraic framework to study and characterize the redundant forms of a biochemical network with a library of constrained, rational number, equivalent and redundant/degenerate matrices for the 𝕏^th^-biochemical network (CRaTER𝕏). These matrices form a many-one map with the stoichiometry number matrix that they are derived from, have distinct mathematical properties, are easily parameterized, can be perturbed and can simulate a biochemical network under differing intracellular conditions. The theoretical results are complemented with detailed computational studies of known biochemical pathways. Future studies will endeavour to search for and identify additional parameters to characterize network-redundancy, identify bounds for the numbers of perturbing PRVs and incorporate/import/analyse data from publicly available datasets.

## 7 Detailed proofs (STx)

Detailed proofs to construct and analyse a reactant-dependent algebraic framework

### 3.4.2 Proofs to define and construct a reactant-dependent algebraic framework

**Proof (T1):**

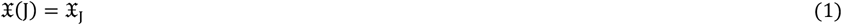

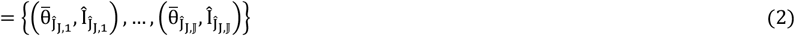

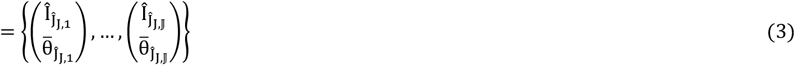

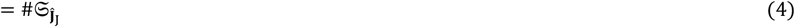

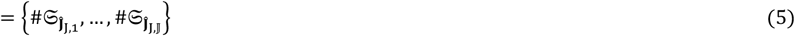

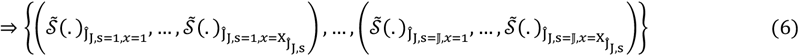

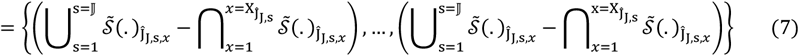

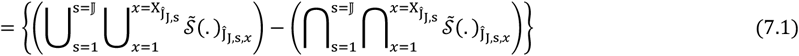

▪

**Proof (C3):**

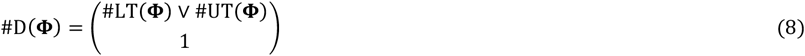

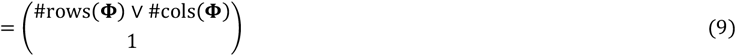

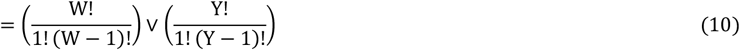

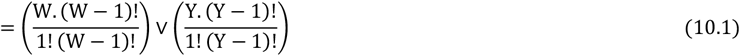

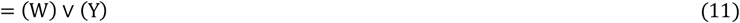

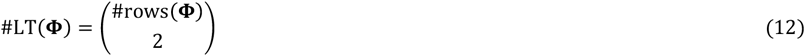

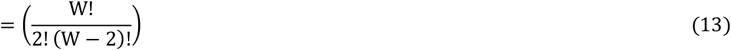

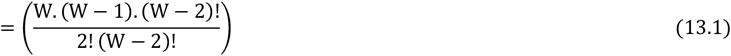

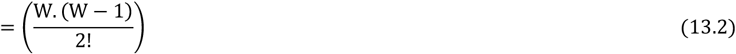

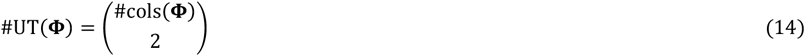

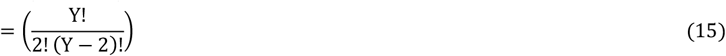

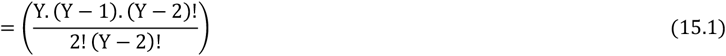

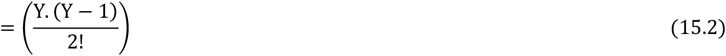

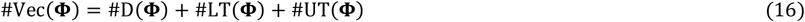

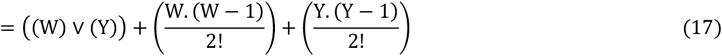

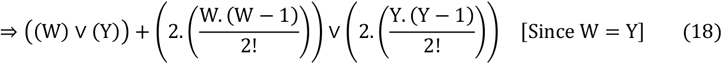

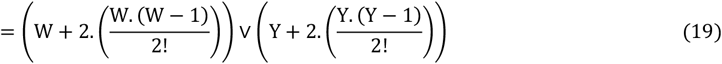

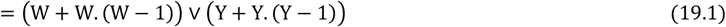

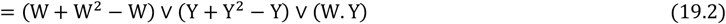

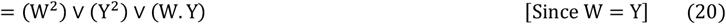

▪

**Proof (C4-C8):**

**C4:**

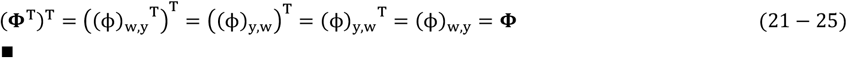

**C5:**

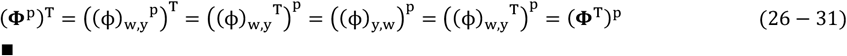

**C6:**

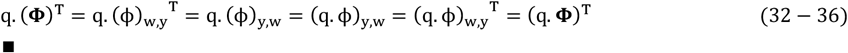

**C7:**

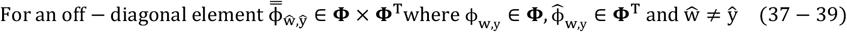

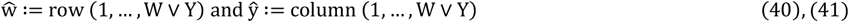

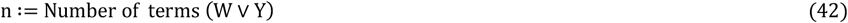

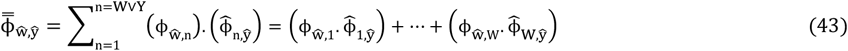

Swapping the row and column indices (transpose) and rewriting Eq. (43)

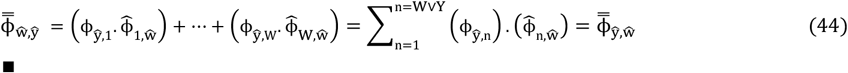

**C8:**

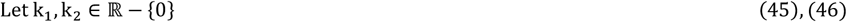

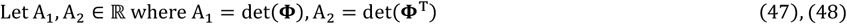

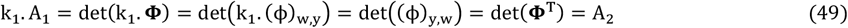

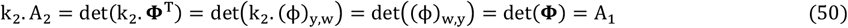

Solving these simultaneously,

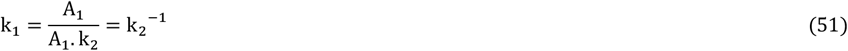

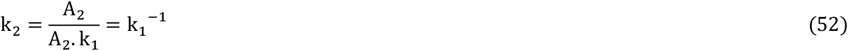

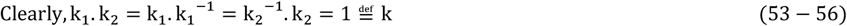

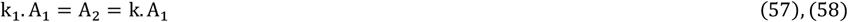

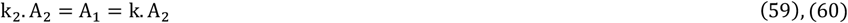

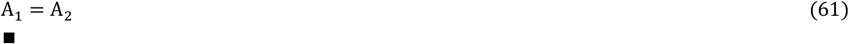

**Proof (T4):**

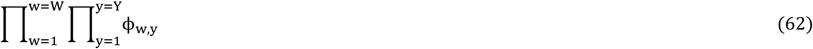

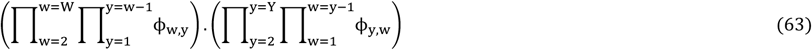

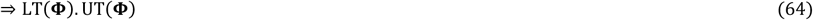

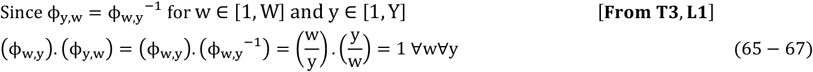

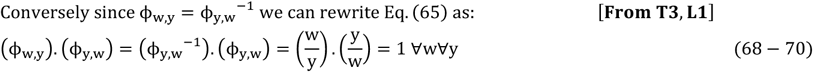

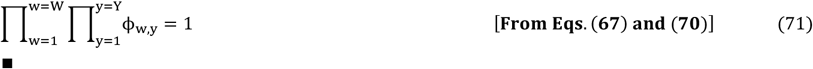

### 3.4.2 Proofs of describe the properties of the redundant forms or equivalent rational number matrices for the 𝕩^th^-biochemical network via the computed null space

**Proof (T7):**

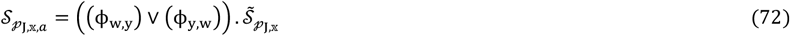

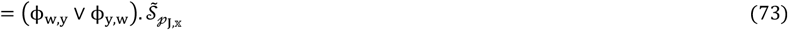

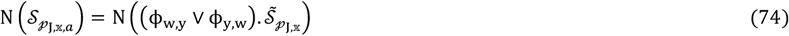

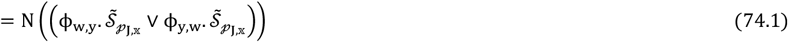

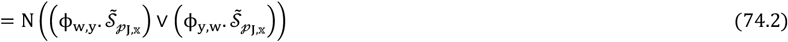

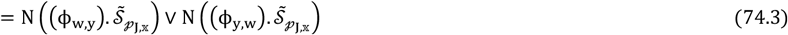

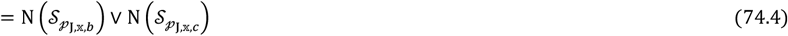

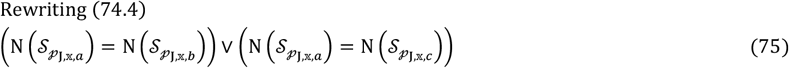

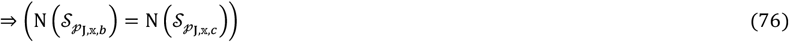

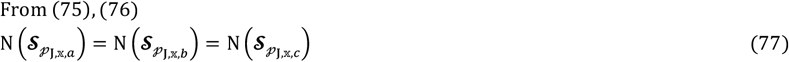

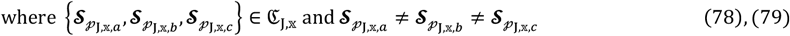

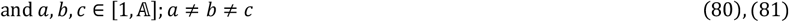

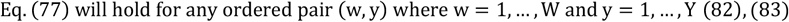

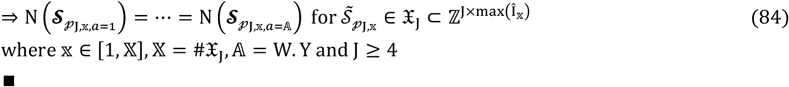

**Proof (T8):** By contradiction

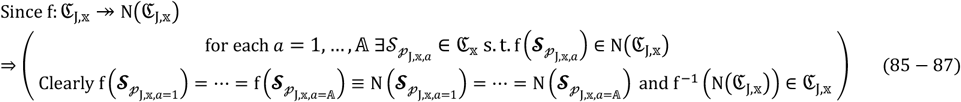

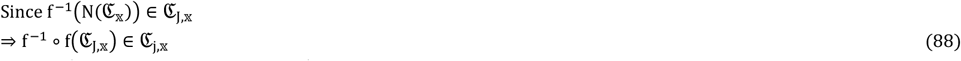

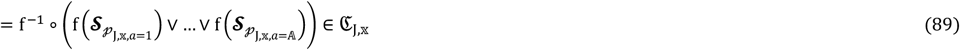

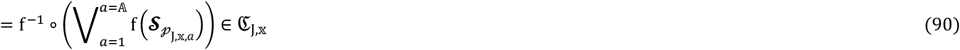

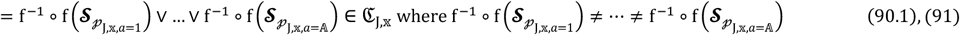

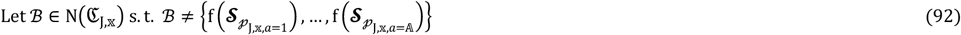

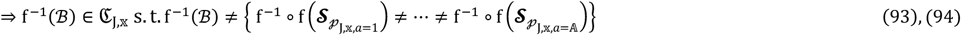

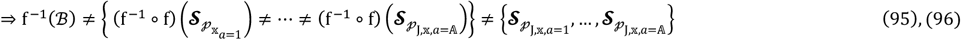

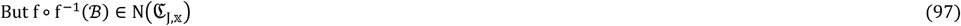

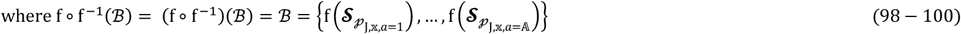

Clearly Eq. (100) contradicts Eq. (92)

▪

### 3.4.2 Proofs to describe the algebraic properties of the redundant forms or equivalent rational number matrices for the 𝕩^th^-biochemical network

**Proof (T9):**

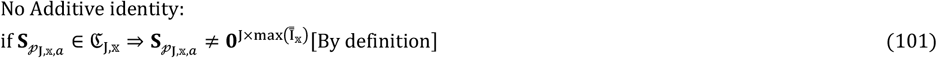

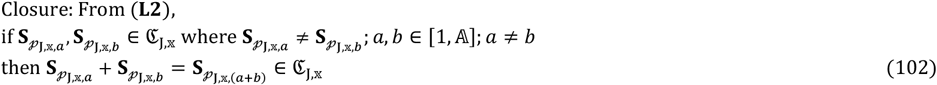

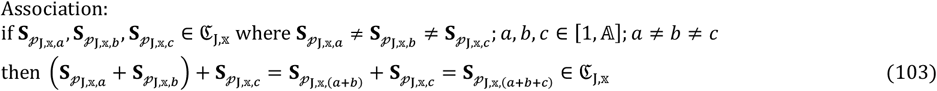

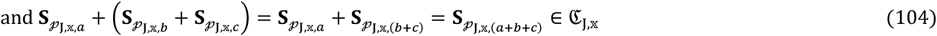

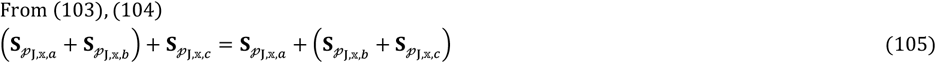

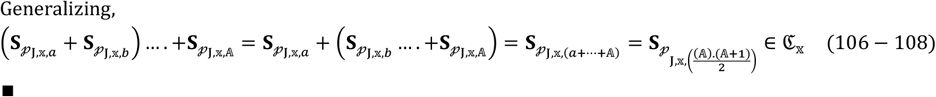

## Acknowledgements

This study was supported, in part by an extramural grant from the Science and Engineering Research Board (SERB), Department of Science and Technology, Government of India under the Mathematical Research Impact-Centric Support (MATRICS) scheme to SK (MTR/2021/000290).

## Author Contributions

SK, designed the study; defined, formulated, developed and wrote proofs for the theorems, lemmas and corollaries; developed the assessment metrics and algorithms; conducted numerical studies and collated data; wrote all necessary code and manuscript.

